# Tumor-Derived Polyamines Initiate Fat Wasting in Cancer Cachexia

**DOI:** 10.64898/2025.12.01.691597

**Authors:** Matias Fabregat, Rachel J Fenske, Julia K Hansen, Cameron J. Kaminsky, Jevin Lortie, Alexander Nassar, Kayla Kressin, Annie Jen, Lucia Cilloni, Katherine Overmeyer, Caroline M Alexander, John W Garrett, Perry J Pickhardt, Joshua J Coon, Costas A Lyssiotis, Marina Pasca di Magliano, Lingjun Li, Adam J Kuchnia, Andrea Galmozzi

## Abstract

Cancer-associated cachexia (CC) is a fatal metabolic condition characterized by progressive loss of fat and muscle mass, yet its early molecular drivers remain poorly defined. Here, we identify a polyamine-dependent tumor-adipose crosstalk that triggers adipocyte lipolysis and fat wasting during the pre-cachexia stage, preceding systemic inflammation and muscle atrophy. Cancer-derived polyamines are enriched in extracellular vesicles and promote lipid mobilization via eIF5A hypusination, independent of adrenergic signaling. In preclinical models, polyamine accumulation associates with early fat loss and elevated circulating fatty acids. Clinically, automated CT imaging of newly diagnosed pancreatic cancer patients reveals increased adipose density, reflecting lipolysis, that correlates with circulating polyamine levels and predicts poor survival. These findings support polyamine metabolism as a mechanistic driver and candidate biomarker of early cachexia, providing a framework for early detection and targeted intervention.

## INTRODUCTION

Cancer-associated cachexia (CC) is a complex metabolic syndrome characterized by involuntary loss of body weight due to accelerated depletion of fat and skeletal muscle (*1-3*). Affecting up to 80% of advanced cancer patients, CC is a major cause of cancer-related mortality (*4-6*). Despite its clinical significance, there are no FDA-approved treatments, and current interventions such as nutritional support and appetite stimulants offer only limited, palliative benefits (*7-16*). The disease progresses through pre-cachexia, cachexia, and refractory stages, with early metabolic changes including heightened adipocyte lipolysis and insulin resistance followed by systemic lipid toxicity, inflammation, and muscle catabolism (*17-21*). Emerging evidence suggests that adipose tissue dysfunction often precedes muscle wasting and serves as a predictor of survival (*22-26*), yet the molecular drivers of adipocyte remodeling remain unknown.

Here, we identify a polyamine-dependent, tumor-adipose metabolic crosstalk that occurs early in pre-cachexia and selectively promotes adipocyte lipolysis prior to other well-characterized systemic changes. Furthermore, we demonstrate that in patients newly diagnosed with early-stage pancreatic cancer, changes in adipose tissue lipolysis can be detected as early as at diagnosis via CT imaging and that such changes correlate with circulating levels of polyamines, establishing a clinically actionable platform for early identification of patients at high risk for cachexia.

## RESULTS

### Cancer-derived polyamines promote adipocyte lipolysis

To model adipocyte dysfunction induced by cancer cells, we adapted and optimized a cell-based system to measure lipid mobilization in primary white adipocytes differentiated *in vitro* and exposed for 24 hours to the conditioned media (CM) of cancer cells derived from different tumor types (Fig. 1A). Mirroring the clinical incidence of cachexia in patients, we found that the CM of cancer cells derived from highly-cachectic tumors, such as pancreatic ductal adenocarcinoma (PDAC), Head & Neck (HNC), colon, and non-small cell lung (NSCLC) cancers, showed a significantly greater induction of lipolysis than the CM of cancer cells derived from tumor types with lower incidence of cachexia, such as breast cancers (Fig. 1, B and C). Notably, no detectable changes in the expression of *Tnfa* or *Ucp1* were observed at the same time point (Fig. 1D), hinting that increased lipolysis precedes the inflammatory response and the beiging of white adipocytes, two well-known events that characterize adipocyte dysfunction in CC (*22, 23*).

**Figure 1.**
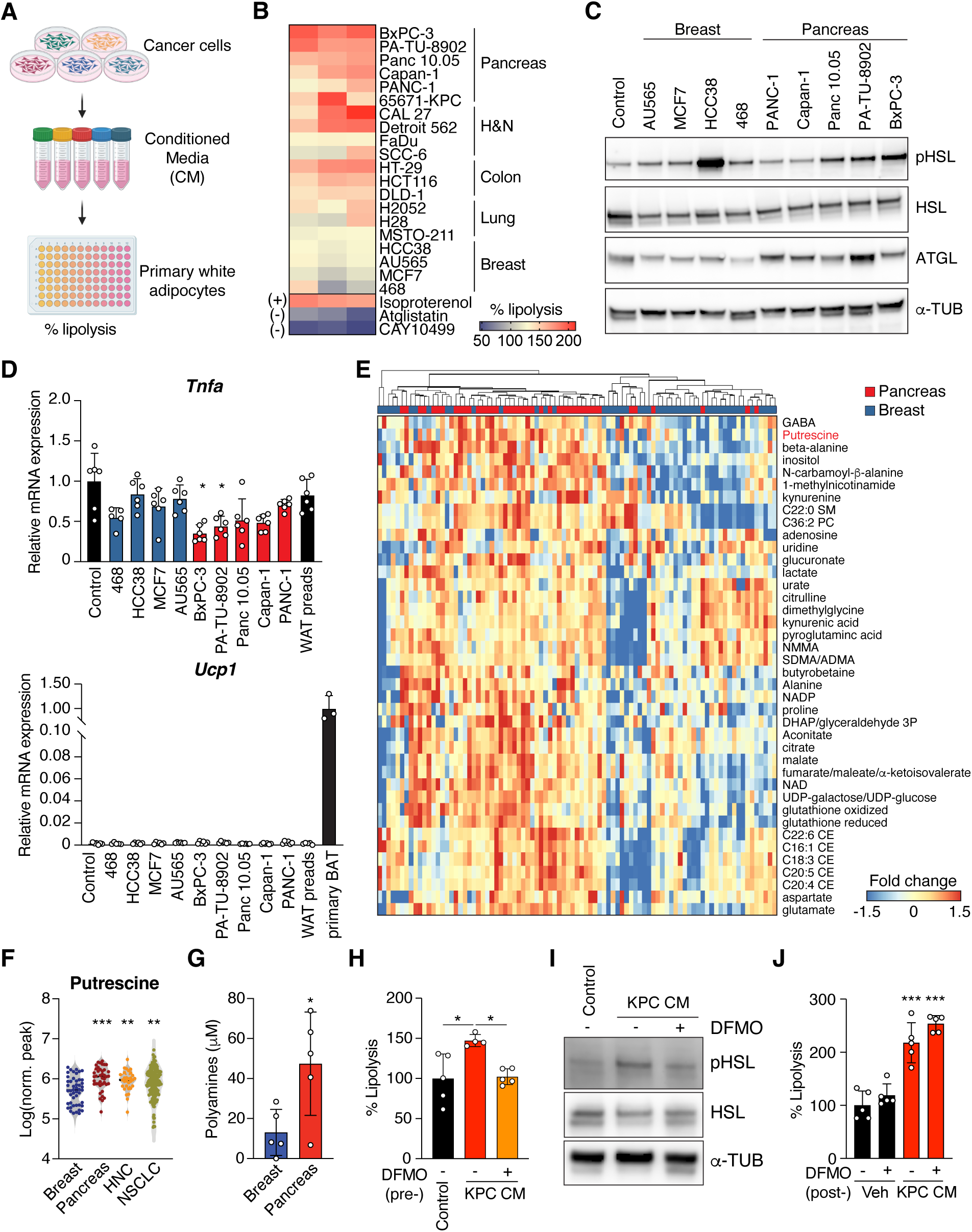
Adipocyte lipolysis is induced in response to cancer-secreted factors. (**A**) Experimental design to interrogate tumor-adipose crosstalk. (**B**) Lipolytic activity in adipocytes exposed for 24 hours to the CM of cancer cells obtained from different tumor types measured as normalized glycerol release into the media. Columns represent the average of three independent experiments. Each experiment was run with n = 3 or more. Isoproterenol was used as a positive control for induced lipolysis, whereas atglistatin and CAY10499, two inhibitors of ATGL and HSL, respectively, were used as negative controls. (**C**) Levels of lipolytic enzymes HSL, as well as its active phosphorylated form pHSL, and ATGL in adipocytes exposed to the CM of pancreatic and breast cancer cells for 24 hours. (**D**) Levels of *Ucp1* and *Tnfa* mRNA in white adipocytes exposed to CM of pancreatic and breast cancer cells for 24 hours (n = 3-6 per condition). (**E**) Relative abundance of metabolites differentially enriched between pancreatic and breast cancer cells profiled in Li et al., 2019 (*29*). (**F**) Intracellular putrescine levels in different pro-cachectic cancer cells compared to non-cachectic cancer cells. Data from Li et al., 2019 (*29*). (**G**) Polyamine quantification in the conditioned media of human breast and pancreatic cancer cell lines used in (B) (n = 5 per group). (**H**) Inhibition of polyamine synthesis with 1 mM DMFO blocks the pro-lipolytic effects of KPC CM (n = 4-5 per condition). (**I**) Protein levels of HSL and pHSL in adipocytes exposed to control media or the CM from KPC treated with either vehicle or DFMO. (**J**) DFMO does not prevent the KPC CM-induced lipolysis if given directly to primary adipocytes. Data shown as mean ± SD. * p<0.05, ** p<0.01, *** p<0.001 by one-way ANOVA with Tukey’s post-test (D, F, I, and J) or unpaired t test (G).

To identify potential factors mediating the effects of pro-cachectic CMs (i.e., PDAC) compared to non-cachectic CMs (i.e., breast), we leveraged the multi-omics dataset of the Cancer Cell Line Encyclopedia (CCLE) (*27-29*) and identified several metabolites differentially abundant between PDAC and breast cancer cell lines (Fig. 1E). Amongst those significantly elevated in PDAC cells, we observed that putrescine levels were also consistently higher in cell lines derived from other highly cachectic cancers, including H&N and NSCLC, compared to less cachectic, breast cancer cells (Fig. 1F). Polyamines are polycationic molecules known for key regulatory roles of cell growth, proliferation, and other biological processes (*30*). Polyamine crosstalk with oncogenic pathways has been widely reported (*31*) and so is the dysregulation of polyamine metabolism in cancer (*32-34*). Interestingly, the expression levels of several proteins involved in the polyamine biosynthetic pathway, including Ornithine Decarboxylase (ODC1), the first and rate-limiting enzyme in polyamine biosynthesis, spermine oxidase (SMOX), and spermidine/spermine N1-acetyltransferase 1 (SAT1), were also significantly higher in PDAC, HNC, and NSCLC cells compared to breast cancer cells (Fig. S1, A and B), whereas negative regulators of polyamine biosynthesis, such as ornithine decarboxylase antizyme 3 (OAZ3) followed the opposite trend (Fig. S1, A and B). Consistent with the metabolite profiling of the CCLE, we found that polyamine concentrations were also significantly higher in the conditioned media of pancreatic cancer cell lines compared to the CM from breast cancer lines (Fig. 1G).

Prompted by these results, we sought to determine the extent to which polyamines contribute to the pro-lipolytic activity of pancreatic cancer cells’ CM. To this end, we treated 65671-KPC cells (hereafter referred to as KPC cells) with either vehicle or the selective, FDA-approved ODC1 inhibitor DFMO (Difluoromethylornithine, 1 mM), and found that inhibition of polyamine synthesis in cancer cells completely abolished the pro-lipolytic effects of their conditioned media (Fig. 1H and I). Importantly, the addition of DFMO to control media or KPC CM post-collection had no impact on adipocyte lipolysis (Fig. 1J), indicating that DFMO prevents the induction of adipocyte lipolysis by blocking polyamine synthesis in cancer cells, rather than through a direct effect on adipocytes. Consistent with these findings, DFMO treatment similarly suppressed the pro-lipolytic activity of conditioned media derived from human pancreatic cancer cell lines BxPC-3 and PA-TU-8902 (Fig. S1C). Finally, DFMO at 1 mM did not significantly impact pancreatic cancer cell viability (Fig. S1D), ruling out the possibility that the loss of lipolytic activity in the conditioned media obtained from DFMO-treated cancer cells was due to cell death. Collectively, these results indicate that cancer-derived polyamines mediate, at least in part, the effects exerted by pancreatic cancer cells on adipocytes.

### Cancer-derived polyamines promote adipocyte lipolysis via eIF5A hypusination

A recent study showed that polyamines may act as ligands for the β_3_-adrenergic receptor (*35*) (Fig. S2A). We replicated this observation, showing that treatment of white adipocytes with two polyamine species showed species-specific, dose-dependent activation of lipolysis (Fig. S2B). Inhibition of the adrenergic signaling with propranolol, an FDA-approved β-blocker, prevented the effects of spermidine on lipolysis (Fig. S2, C and D), confirming the ability of spermidine to act as an agonist for the β_3_-adrenergic receptor. However, unlike spermidine treatment alone, co-administration of propranolol with pancreatic cancer cells’ CM, which contain high levels of polyamines (Fig. 1G) and whose effects were reversed by inhibition of polyamine synthesis with DFMO (Fig. 1H and S1E), failed to suppress lipolysis (Fig. S2, E to G). Collectively, these results demonstrate that cancer-derived, polyamine-dependent induction of adipocyte lipolysis occurs through mechanisms distinct from adrenergic signaling.

Besides acting as a ligand for the β_3_-adrenoceptors, spermidine has also been shown to play a critical role in the hypusination of the eukaryotic translation initiation factor 5A (eIF5A) (Fig. 2A), a 2-step activating post-translational modification catalyzed by the enzymes deoxyhypusine synthase (DHPS) and deoxyhypusine hydroxylase (DOHH) (*36*) (Fig. 2A). Therefore, we tested whether cancer-derived polyamines could influence this process. Notably, we observed a striking increase in eIF5A hypusination in white adipocytes upon exposure to KPC CM (Fig. 2B). In parallel, we also detected a significantly higher expression of *Dohh*, whereas the expression of *Dhps* remained unchanged (Fig. S3A). To test whether hypusination of eIF5A by cancer-derived polyamines is a required step to promote lipid mobilization in adipocytes, we exposed cultured adipocytes to KPC CM supplemented with either vehicle or 20 μM GC7 (N(1)-guanyl-1,7-diaminoheptane), a selective inhibitor of the hypusinating enzyme DHPS. Notably, in the presence of GC7, the hyper-hypusination of eIF5A induced by KPC CM was completely prevented, and so was the induction of lipolysis (Fig. 2, C and D), indicating that polyamine-dependent hypusination of eIF5A is required for the induction of adipocyte lipolysis in response to cancer cells conditioned media.

**Figure 2.**
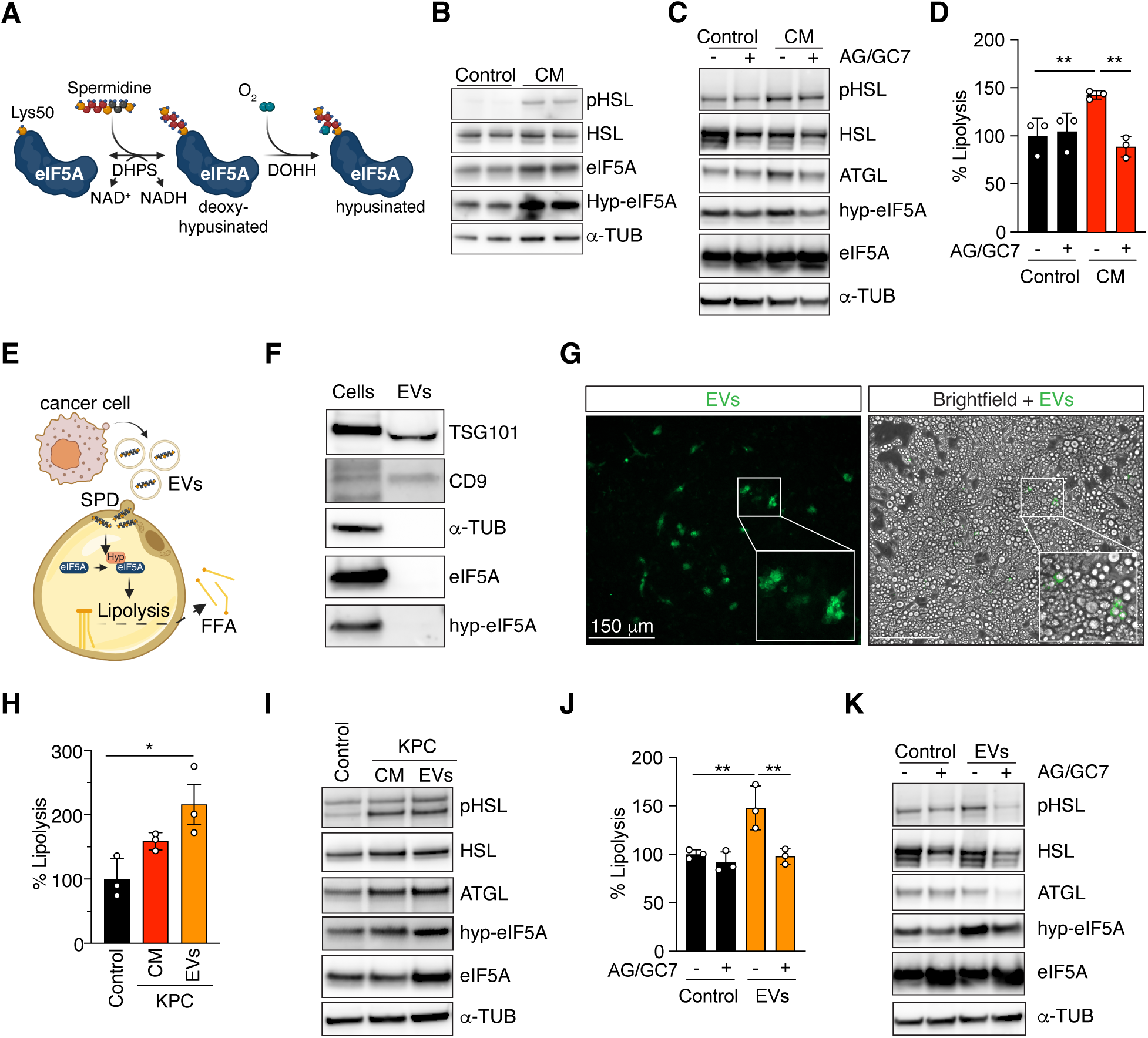
Polyamine-dependent hypusination of eIF5A is required for the induction of adipocyte lipolysis. (**A**) Spermidine is the substrate for the 2-step post-translational modification of eIF5A. (**B**) KPC CM induces hypusination of eIF5A in adipocytes. (**C**, **D**) Inhibition of eIF5A hypusination with AG/GC7 (AG=aminoguanidine, a GC7 stabilizer) prevents hypusination and lipolysis in adipocytes exposed to KPC CM (n = 3). (**E**) Schematic representation of cancer EVs’ delivery of polyamines to the adipocytes. (**F**) EV markers TSG101 and CD9, but not hypusine or eIF5A, can be detected in purified EVs. (**G**) Representative images of fluorescently labeled EVs taken up by adipocytes 24 hours post-incubation. (**H**, **I**) KCP EVs promote lipolysis and eIF5A hypusination in adipocytes to a similar extent of the complete KPC CM (n = 3). (**J**, **K**) Inhibition of eIF5A hypusination with AG/GC7 prevents hypusination and lipolysis induced by KPC’s EVs (n = 3). Data shown as mean ± SD. * p<0.05, ** p<0.01, *** p<0.001, 1-way ANOVA-Tukey’s post-test or unpaired t test (C).

We then wonder how cancer-derived polyamines can selectively act via eIF5A hypusination rather than by activating the adrenergic receptor. We hypothesized that cancer-derived polyamines may not be freely secreted into the extracellular environment but may be “packaged” into extracellular vesicles (EVs), thus bypassing the adrenergic receptor on the plasma membrane. Upon uptake by the adipocytes, polyamines can then be released into the cytosol and serve as readily available substrates for the hypusinating enzymes (Fig. 2E). To test this model, we purified EVs from the CM of KPC cells (Fig. 2F and S3B) and treated mature white adipocytes with these particles for 24 hours. Fluorescently labeled EVs were efficiently internalized by adipocytes (Fig. 2G) and, mirroring KPC CM, were both able and sufficient to promote adipocyte lipolysis and induce eIF5A hypusination (Fig. 2, H and I). Importantly, inhibition of eIF5A hypusination via co-treatment with GC7 completely abolished the lipid-mobilizing effects of KPC EVs (Fig. 2, J and K), demonstrating that KPC EVs promote lipolysis through hypusination of eIF5A.

To determine whether this newly identified mechanism also regulates lipolysis in adipose tissue, we purified EVs from the CM of KPC cells and treated inguinal white adipose tissue (iWAT) explants *ex vivo* with either EVs or EV-depleted CM (Fig. S3C). Consistent with our observations in cultured adipocytes, KPC-derived EVs robustly stimulated lipolysis in tissue explants (Fig. S3D), and these effects were fully reversed either by treating KPC cells with DFMO (Fig. S3E) or by co-incubating adipose explants with the hypusination inhibitor GC7 (Fig. S3F). Strickingly, EV-depleted CM failed to induce lipolysis (Fig. S3D), indicating that the pro-lipolytic activity of KPC CM is mediated primarily, if not exclusively, by polyamine-enriched EVs released from cancer cells.

Finally, neither eIF5A nor its hypusinated form was detected in KPC-derived EVs (Fig. 2F), thereby excluding the possibility that EVs promote lipolysis through direct transfer of eIF5A protein rather than through delivery of its metabolic activators.

### Cancer-induced fat wasting precedes muscle loss and correlates with circulating polyamines *in vivo*

To investigate how cancer-derived polyamines contribute to cachexia *in vivo*, we used a syngeneic mouse model in which KPC cells (5 × 10^5^ cells/animal) were injected subcutaneously into the flanks of FVB female mice. In this model, tumors become palpable about seven days after implantation and typically reach humane endpoints within five weeks (Fig. 3A). Mice were monitored every two days for body weight, tumor volume, and food consumption, and weekly for body composition and grip strength. By four weeks post-implantation, overall body weight and food intake were unchanged compared with controls (Fig. 3, B and C), indicating that overt cachexia or anorexia had not yet developed. However, body composition analysis revealed a significant reduction in fat mass (Fig. 3, D and E, and Fig. S4A). Conversely, tumor-adjusted lean mass remained stable at this stage (Fig. 3, D and E), indicating that lipid stores are affected earlier than skeletal muscle during disease progression. Supporting this interpretation, expression of classical muscle atrophy markers, including MuRF1 (*Trim63*) and Atrogin-1 (*Fbxo32*), remained unchanged at the four weeks timepoint (Fig. S4B), and muscle weakness, as measured by grip strength, appeared only later, around five weeks post-implantation (Fig. S4C).

**Figure 3.**
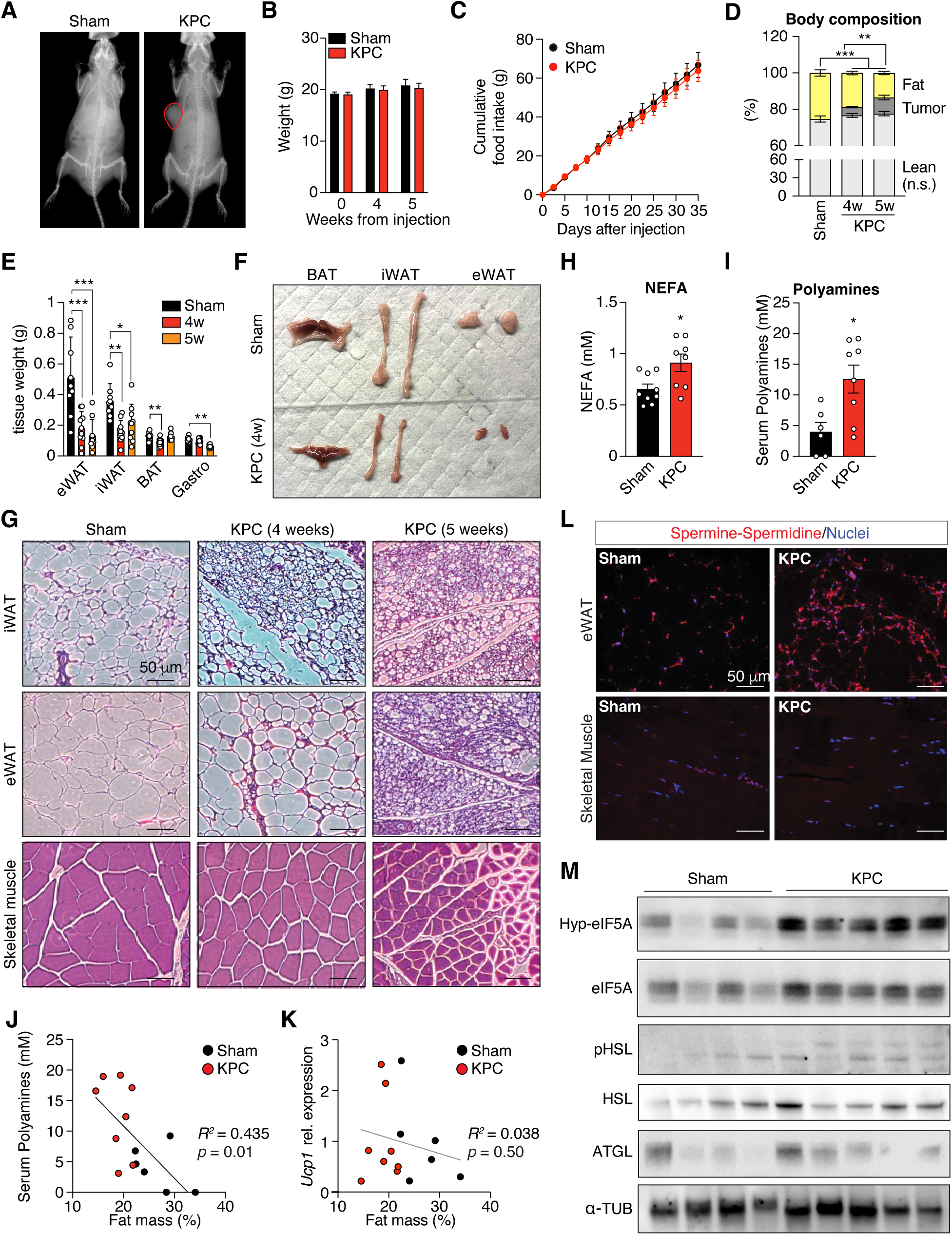
Adipose tissue wasting precedes muscle loss in KPC tumor-bearing mice. (**A**) DEXA scan of sham and tumor-bearing mice four weeks after cancer cell injection. (**B**) Body weight progression of sham and tumor-bearing mice (n = 9 sham; 20 KPC four weeks, 10 for five weeks). (**C**) Cumulative food consumption per mouse in sham and tumor-bearing mice over the course of five weeks (n = 9 sham; 20 KPC, 10 after day 28). (**D**) Body mass composition of sham and tumor-bearing mice reveals loss of fat mass precedes muscle waste (n = 9 sham; 20 KPC, 10 after day 28). (**E**) Adipose and muscle weights in sham and tumor-bearing mice at four and five weeks post-injection (n = 9 sham, 10 KPC four weeks, 10 KPC five weeks). (**F**) Gross appearance of adipose depots in sham and tumor-bearing mice at four weeks. (**G**) H&E staining of iWAT, eWAT, and gastrocnemius muscle in sham and tumor-bearing mice at four and five weeks. (**H**, **I**) Circulating NEFA and polyamine levels are elevated in tumor-bearing mice at four weeks (n = 6-9 sham, 8 KPC). (**J**) Circulating levels of polyamines correlate with adipose tissue mass (n = 6 sham, 8 KPC). (**K**) Ucp1 expression in iWAT does not correlate with adipose tissue mass (n = 6 sham, 8 KPC). (**L**) IF staining of polyamines in eWAT and skeletal muscle of sham and KPC mice at four weeks reveals selective accumulation of polyamines in adipose but not in muscle. (**M**) Protein levels of lipolytic enzymes and hypusinated eIF5A in eWAT of sham and KPC tumor-bearing mice at four weeks. Data shown as mean ± SEM. *p<0.05, **p<0.01, ***p<0.001, 1-way ANOVA with Dunnett’s post-test (D, E) or unpaired t test (H, I).

Gross morphology and histological analysis confirmed these observations. In fact, all the adipose depots analyzed, including inguinal (iWAT), epididymal (eWAT), and dermal (dWAT) white adipose tissues, exhibited reduced tissue weight, smaller adipocytes, and increased fibrosis by four weeks (Fig. 3, F and G, and Fig. S4, D to F), while skeletal muscle morphology remained unaffected until later stages (Fig. 3G and Fig. S4D). Collectively, these results indicate that fat wasting precedes any detectable muscle loss. Consistent with early fat mobilization, circulating non-esterified fatty acids (NEFA) were significantly elevated at four weeks (Fig. 3H), reflecting active lipolysis. Importantly, paralleling the increased circulating polyamines observed in cachectic cancer patients (*32-34, 37-40*), tumor-bearing mice also showed higher serum polyamine levels than sham controls (Fig. 3I). Furthermore, serum polyamine concentrations inversely correlated with fat mass (Fig. 3J), hinting at a possible causal link between cancer-derived polyamines and adipose tissue catabolism.

At this time point, there were no indications of tissue inflammation or white adipose tissue beiging, two hallmarks of more advanced stages of cachexia (*22, 23*), for no significant changes were observed in iWAT and eWAT inflammatory (*Tnfa*) and thermogenic (*Ucp1*) markers (Fig. S4G). Accordingly, Ucp1 expression in iWAT showed no correlation with fat mass (Fig. 3K).

Finally, at four weeks post-implantation, elevated polyamine levels were detected not only in circulation but also within the adipose tissue itself (Fig. 3L). In eWAT, this local accumulation was accompanied by a marked increase in eIF5A hypusination as well as higher levels of phospho-HSL and ATGL (Fig. 3M), indicating activation of polyamine-associated signaling within the tissue. In stark contrast, skeletal muscle showed no comparable rise in polyamine content at this stage (Fig. 3L), reinforcing that the adipose tissue serves as the earliest and most responsive target of cancer-derived polyamine signaling. Together, these results support a model where polyamine-driven lipolysis represents an inflammation- and beiging-independent event that contributes to adipose tissue wasting during the early stages of cachexia progression.

### Automated CT Imaging Identifies Early Adipose Remodeling in Pre-Cachexia

Despite decades of research, no reliable method exists to diagnose cachexia in its early or pre-cachectic stages, when metabolic alterations are underway but precede overt weight loss (*41-44*). Conventional clinical metrics such as body mass index (BMI) fail to detect these patients early enough for effective intervention (*41-44*). Routine staging computed tomography (CT) scans, universally obtained at cancer diagnosis, offer an underused opportunity to detect early metabolic dysfunction (*43, 45*). These images contain quantitative information on adipose and skeletal muscle composition that can reveal subtle changes in lipid content and fibrosis (*43, 45-47*).

To leverage this potential, we implemented an automated CT body composition analysis pipeline that extracts reproducible metrics from standard-of-care scans. The pipeline quantifies total (TAT), visceral (VAT), subcutaneous (SAT), and intermuscular (IMAT) adipose tissue areas and densities, as well as skeletal muscle area and density (Fig. 4A). This automated approach minimizes user bias, enhances reproducibility, and is validated across both contrast-enhanced and non-contrast CT datasets (*48-52*). Using this computational framework, we retrospectively analyzed diagnostic CT scans from 39 patients newly diagnosed with PDAC (Supplementary Table S1). Consistent with prior studies (*41-44*), BMI and skeletal muscle area did not predict two-year survival (Fig. 4, B and C). In contrast, patients who died within two years of diagnosis exhibited significantly lower TAT and VAT areas (Fig. 4, D and E). SAT and IMAT areas also showed similar trends but did not reach statistical significance when analyzed independently (Fig. 4E). Even more prognostic was the adipose tissue density, a CT-derived marker of lipid mobilization (*53*). Patients with survival under two years exhibited higher TAT, VAT, SAT, and IMAT densities, but not muscle density, consistent with enhanced adipocyte lipolysis and adipose tissue remodeling (Fig. 4F). Importantly, CT-derived metrics were significantly associated with two-year survival independent of cancer stage, as none of the CT measures differed across stages (Fig. S5A). Circulating non-esterified fatty acids (NEFAs) also trended upward in patients with poor survival (Fig. 4G), supporting increased lipolytic activity in these subjects. Conversely, at diagnosis, other known mediators of cachexia, including GDF15, IL-6, and markers of muscle damage such as creatine kinase (CKM), showed no association with two-year survival (Fig. S5B). These results indicate that adipose tissue remodeling precedes activation of canonical cachexia pathways, can be detected as early as at diagnosis, and is associated with poor outcomes.

**Figure 4.**
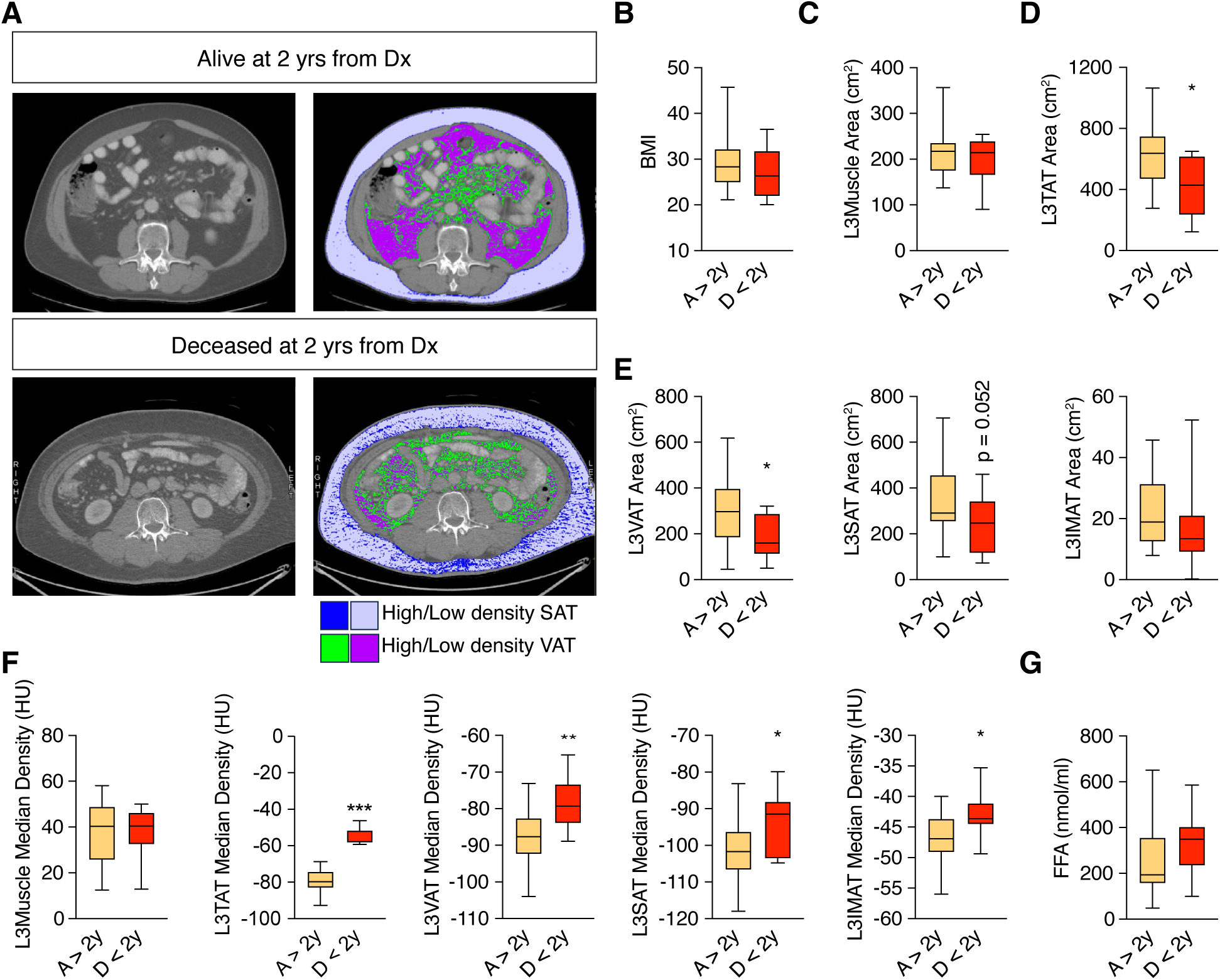
CT imaging detection of early adipose remodeling in PDAC patients. (**A**) Representative diagnostic CT scans and automated analysis of lean and fat mass area and density in patients newly diagnosed with pancreatic ductal adenocarcinoma. Lavender and violet define SAT and VAT areas, respectively, while blue and green represent high density regions within SAT and VAT. (**B**, **C**) Body mass index (BMI) and skeletal muscle area at diagnosis are not predictive of survival (n = 30 A > 2y, 9 D < 2y). (**D**, **E**) Total (TAT), visceral (VAT), subcutaneous (SAT), and intermuscular (IMAT) adipose tissue areas are already reduced at diagnosis in patients with poor outcomes (n = 30 A > 2y, 9 D < 2y). (**F**) Adipose tissue, but not skeletal muscle, density is significantly elevated at diagnosis in PDAC patients with poor outcomes and is associated with two-year survival (n = 30 A > 2y, 9 D < 2y). (**G**) Circulating free fatty acid trend upward in PDAC patients with poor survival (n = 30 A > 2y, 9 D < 2y). A, alive; D, dead. Data shown as median + min/max intervals. *p<0.05, ***p<0.001, unpaired t test.

### Polyamine accumulation links early tumor progression to adipose remodeling

In patients with cancer, circulating polyamine levels peak during the early stages of tumor progression (stages I-II) (*38, 40*). Consistent with this observation, serum polyamine concentrations at diagnosis were significantly higher in early-stage (I-II) PDAC patients compared with those diagnosed at advanced stages (III-IV) (Fig. 5A). In contrast, circulating levels of GDF15, IL-6, and CKM did not differ significantly between early- and late-stage patients (Fig. 5B). Notably, within the early-stage cohort, polyamines, and in particular putrescine species, positively correlate with IMAT area, a marker of fat infiltration known to precede weight loss and predict cachexia severity (*54-56*), as well as with VAT and SAT density (Fig. 5C). These associations suggest a mechanistic link between polyamine metabolism and adipose tissue remodeling in patients with pancreatic cancer. Prompted by these findings, we next examined whether elevated serum polyamines could predict survival in early-stage patients. The cohort was stratified into “high PA” or “low PA” groups based on median serum polyamine levels and survival curves were generated for both groups. Strikingly, patients with high serum polyamines at diagnosis exhibited significantly reduced 5-year survival (Fig. 5D). Collectively, these results support a model in which polyamine-driven lipolysis contributes to early metabolic reprogramming, leading to measurable increases in adipose tissue density on CT imaging. This metabolic-imaging axis may therefore serve as an early, noninvasive biomarker of cachexia onset.

**Figure 5.**
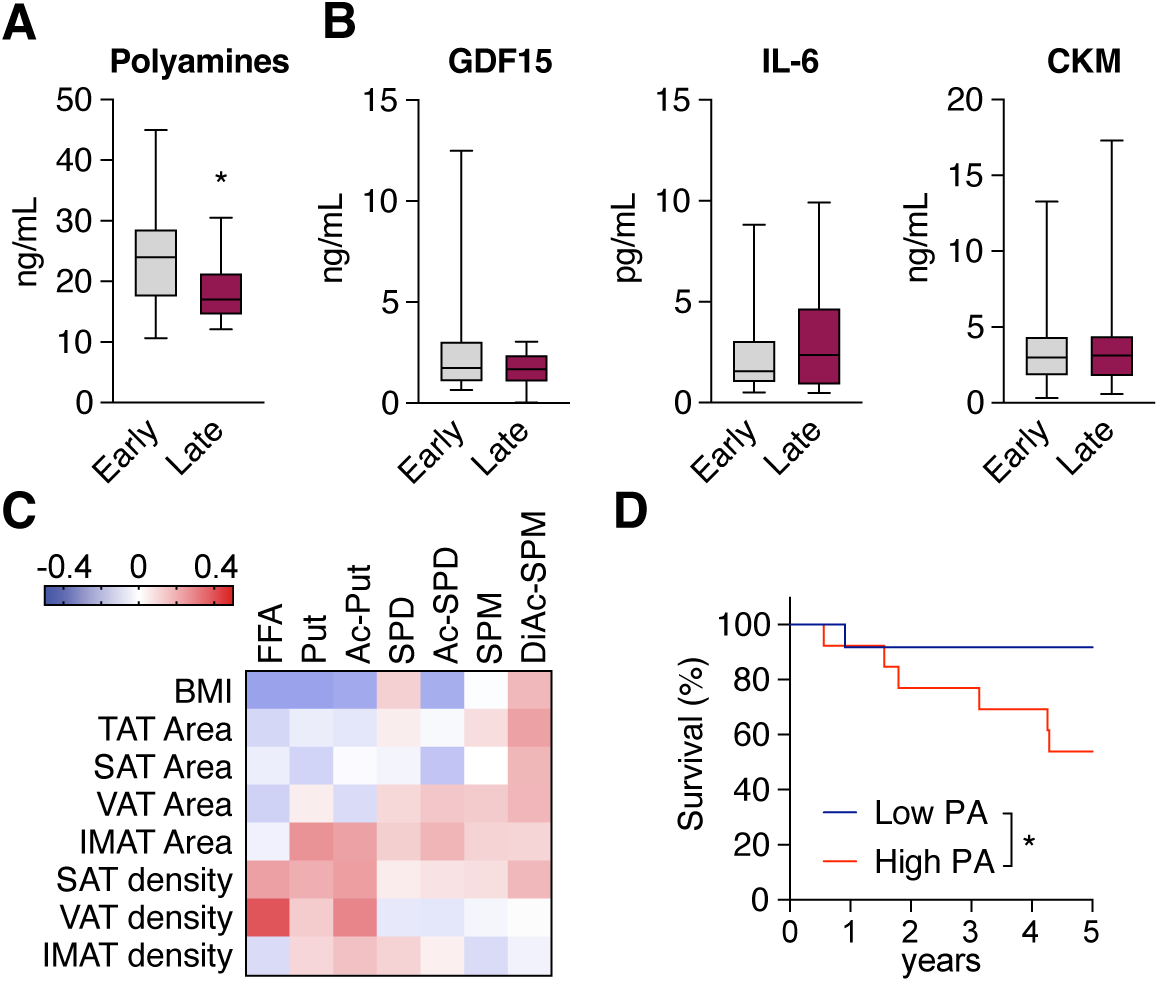
Circulating polyamines correlate with adipose tissue metrics in early stage PDAC patients. (**A**) Total levels of circulating polyamines are significantly elevated in early-stage (I and II) PDAC patients compared to late-stage (III and IV) PDAC patients (n = 25 Early, 12 Late). (**B**) Circulating levels of GDF15, IL-6, and CKM are not different between early- and late-stage PDAC patients (n = 25 Early, 12 Late). (**C**) Correlation plot of circulating polyamine and free fatty acids with BMI and CT metrics shows direct correlation between putrescine and free fatty acids with IMAT area and SAT and VAT density (Put = putrescine; Ac-Put = acetyl-putrescine; SPD = spermidine; Ac-SPD = acetyl-spermidine; SPM = spermine; DiAc-SPM = diacetyl-spermine). (**D**) Five-year survival curves in patients with stage 1 and II pancreatic cancer and circulating levels of polyamines below (Low PA) or above (High PA) the cohort median. Data shown as median + min/max intervals. *p<0.05, unpaired t (A) or Logrank (Mantel-Cox) test.

## DISCUSSION

Our findings define a tumor-adipose signaling axis in which cancer-derived polyamines act as early inducers of adipocyte lipolysis. Polyamines, enriched in tumor-secreted extracellular vesicles (EVs), are internalized by white adipocytes and trigger eIF5A hypusination, activating lipolytic programs independently of adrenergic signaling. This process mobilizes stored triglycerides, activates ATGL and HSL in white adipose depots, and elevates circulating NEFAs, all before systemic inflammation, beiging, or skeletal muscle atrophy arise, identifying eIF5A hypusination as an early metabolic trigger of cachexia.

Our results are consistent with prior studies showing that lipolysis, rather than adipocyte death, drives adipose wasting in cachexia (*19, 20*). They support a model in which lipid efflux from adipose tissue promotes ectopic lipid deposition (e.g., intermuscular fat deposition), systemic lipotoxicity, and insulin resistance (*57*), while simultaneously activating hypothalamic circuits that amplify glucocorticoid output and further accelerate lipid and protein catabolism (*21, 58*). Thus, in this model, the adipose tissue is not merely consumed by systemic inflammation but serves an early and direct target of tumor-derived metabolites.

Temporally, our data suggest that the polyamine-eIF5A hypusination axis operates upstream of established inflammatory and neuroendocrine mediators of advanced cachexia, including IL-6 (*23*), TNFα (*59*), IL-1 (*60*), PTHrP (*22*), and ZAG (*24*), which fuel systemic inflammation, beiging of white adipose tissue, thermogenesis, and whole-body energy expenditure. Consistent with this interpretation, tumor-bearing mice exhibit selective loss of fat mass, adipose fibrosis, and elevated NEFAs at a stage when total body weight, food intake, lean mass, and markers of inflammation and thermogenesis remain unchanged.

Genetic deletion of ATGL, the rate-limiting enzyme for triglyceride breakdown, preserves adipose mass and protects muscle from apoptosis in tumor-bearing mice (*25*), underscoring that selective inhibition of tumor-induced lipolysis in adipocytes could provide metabolic protection and maintain energy balance. However, the broad tissue distribution of lipolytic enzymes limits the use of pharmacological agents to inhibit ATGL or HSL, as systemic inhibition of lipolysis may cause severe side effects (*61, 62*). Our results identify novel molecular nodes that could be targeted to locally suppress lipolysis, maintain adipose metabolic integrity, and dampen the neuroendocrine-inflammatory drive of systemic catabolism.

Routine staging CT scans performed at diagnosis offer a practical and cost-free opportunity to identify patients with early adipose dysfunction (*43, 45*). Using an automated CT-based body composition pipeline, we demonstrate that adipose density, rather than BMI or skeletal muscle area, is strongly predictive of survival in patients with pancreatic ductal adenocarcinoma. These imaging-derived metrics are independent of nutritional status at the time of scanning, in contrast to circulating NEFAs that fluctuate with feeding state, underscoring their robustness as potential metabolic biomarkers. The correlation between circulating polyamines and adipose remodeling further suggests that tumor-derived polyamines may serve as both mechanistic drivers and measurable indicators of early cachexia. In fact, in patients with early-stage PDAC, we show that polyamines correlate with CT-metrics of enhanced adipose lipolysis, even when BMI, muscle area, and other mediators of cachexia such as IL-6, GDF-15, and CKM remain unaffected.

This correlation between circulating polyamines and imaging signatures suggests that integrating serum and radiomic metrics could enable early, minimally invasive identification of patients at highest metabolic risk. Therapeutic strategies that target this early tumor-adipose crosstalk, combined with imaging and metabolic biomarkers for patient stratification, shift the focus from late-stage, symptom-directed interventions to early detection and prevention of metabolic dysfunction.

More broadly, by integrating molecular, imaging, and clinical data, this work establishes a foundation for a clinically applicable risk-prediction model to identify patients at high risk for cachexia. Targeting polyamine synthesis or eIF5A hypusination may preserve metabolic integrity, improve tolerance to antineoplastic therapy, and ultimately extend survival and quality of life in patients with pancreatic cancer.

## MATERIAL AND METHODS

All experiments were conducted at the University of Wisconsin Madison. Study protocols were approved by the animal care and use committee of the University of Wisconsin Madison School of Medicine and Public Health and conducted under ethical regulations and policies.

### Cancer Cell Line Encyclopedia database

Metabolomics and RNAseq data from the Cancer Cell Line were downloaded from the DepMap Portal website (https://depmap.org/portal/data_page/?tab=allData) from the CCLE2019 files.

Heatmaps of scaled metabolite abundance were generated using pheatmap in R with sample annotation columns representing tissue or cancer type. The heatmaps were color-scaled using palettes from RColorBrewer and ordered hierarchically.

### Laboratory animals

FVB female mice were obtained from Jackson Laboratory and housed under pathogen-free conditions on a 12-hour light-dark cycle with unlimited access to water and food (Chow diet, Teklad #2018). Before any experimental procedures, mice were allowed to acclimate for at least 7 days. For tumor induction, 12-week-old female mice were randomly divided into two experimental groups, anesthetized with isoflurane, and each mouse received a subcutaneous injection of 200 µL of either PBS (Sham) or 5 × 10^5^ 65671-KPC cells in 200 µL of PBS (KPC). Before injection, 65671-KPC cells were dissociated, counted using a hemocytometer (Kova Plastics, #22-270141), and resuspended in PBS supplemented with 0.5% of BSA and 25mM of Glucose at a concentration of 2.5 × 10^6^ viable cells/mL. Mice were monitored every two days for body weight and food consumption and weekly for body composition and grip strength with minimal handling to reduce stress. Body composition analysis was measured using an EchoMRI (EchoMRI-500-A100-A10TM, EchoMRI LLC). Hind-limb grip strength was measured using a grip strength meter (San Diego Instruments). Test was performed by allowing the animal to grasp the grid with both hind limbs and then pulling the animal straight away from the grid until it released the platform. In each session, the average peak force of the top three from five attempts was recorded.

For molecular analysis of tissues, mice were euthanized and tissues collected right after, snap-frozen, and stored at -80°C.

### Histology and Immunostaining

Following euthanasia, aliquots of tissues were collected and fixed in 10% neutral buffered formalin for 24 hours, extensively washed with PBS and stored in 70% ethanol. Tissues were dehydrated, embedded in paraffin and 3mm thick sections stained with hematoxylin and eosin by the UW Experimental Animal Pathology Lab. Images were acquired using an EVOS M5000 Imaging System (Thermo Fisher), and lipid droplet size and muscle fiber area were quantified using ImageJ software.

Antigen retrieval was performed in the R-Universal Epitope Recovery Buffer (Electron Microscopy Sciences, #62719) using the 2100 Retriever heat-induced epitope recovery system (Electron Microscopy Sciences). Immunostaining was performed as previously described (*63*). The primary antibody used for polyamine detection was the rabbit anti-SPD/SPM (1:500, Novus Biological NB1001847). The secondary antibody was Biotin-SP-conjugated AffiniPure goat anti-rabbit (Jackson ImmunoResearch Laboratories, #111-065-003) followed by Alexa Fluor 594-conjugated streptavidin (Jackson ImmunoResearch Laboratories, # 016-580-084). Nuclei were counterstained with DAPI 4′,6-Diamidino-2phenylindole dihydrochloride (Thermo Fisher #62248). Negative controls were performed with no primary antibody. Images were captured on an Axio Observer 5 microscope (Carl Zeiss Microscopy) with identical settings between experimental groups, and analyzed using ImageJ software.

### Serum analysis

Blood collection was performed retro-orbitally post euthanasia. Whole blood was allowed to clot at room temperature for 30 minutes and spun down at 1,500 g for 15 minutes. The supernatant was collected and transferred to a new tube and then spun down for 30 seconds at 18,500 g. Supernatant was then transferred again into a new tube and stored at -80°C until further analysis. Non-esterified fatty acids were quantified using the commercial kits from WAKO Fujifilm, HR Series NEFA-HR (2) Color Reagent A (cat# 999-34691), HR Series NEFA-HR (2) Color Reagent B (cat# 991-34891), HR Series NEFA-HR (2) Solvent A (cat# 995-34791), HR Series NEFA-HR (2) Solvent B (cat# 993-35191), NEFA Linearity Material (cat# 997-76491), NEFA Standard Solution (cat# 276-76491). Total polyamine levels were quantified using the commercial kit from Sigma Aldrich (MAK349).

### *Ex vivo* experiments

All *ex vivo* experiments were performed using inguinal (iWAT) fat explants. For these experiments, 10-12 weeks old C57BL6/J mice were used. Mice were euthanized and fat depots were collected immediately after. Upon dissection and weighing, fat depots were transferred into 10 cm dishes containing 10 mL of prewarmed DMEM. Tissues were minced into approximately 50 mg biopsies. Each biopsy was then moved into one well of a 24-well plate containing 1 mL of prewarmed DMEM and maintained overnight at 37 °C. The following day, media was renewed to FBS-free DMEM supplemented with 2 % fatty acid-free BSA along with the specified stimuli. Fat explants were then incubated for 24hs at 37 °C and glycerol content in the media was measured and normalized to non-stimulated condition. Finally, fat explants were washed with PBS and processed for protein extraction or stored at -80 °C for further analysis.

### 65671-KPC cell culture and injection

65671-KPC cells (*64*) were cultured in DMEM (Thermo Fisher, #11965118) supplemented with 10% heat-inactivated fetal bovine serum (Gemini, #100106) and 1% antibiotic-antimycotic solution (Thermo Fisher, #15240062). The cells were maintained in a humidifier incubator at 37°C with 5 % CO_2_. For expansion, frozen KPC cells were rapidly thawed, centrifuged at 1200g for 5 minutes, and cultured in 10 cm dishes. Upon reaching confluence, cells were dissociated using 0.25 % Trypsin-EDTA buffer (Thermo Fisher, #25300054), centrifuged, and subcultured into 15 cm dishes at a 1:5 split ratio until sufficient cells were obtained for tumor cell injection.

### Conditioned media collection

Cancer cells were cultured in DMEM supplemented with 10 % FBS. At 80 % confluency, media was replaced with DMEM supplemented with 2 % fatty acid-free BSA and cells were allowed to grow for 48 hours. The conditioned media was then collected, spun down for 10 minutes at 300 g, and the supernatant was filtered through a 0.2 μm polyethersulfone filters (MilliporeSigma) into a new tube. Finally, glucose was added back to a final concentration of 25 mM and CM was stored at -80 °C until further use.

### Cell viability assay

Cancer cells were cultured up to 80% of confluence, after which dosing with DFMO, or equal concentration of DMSO (DFMO solvent) was performed (0, 0.25, 0.5, 1, 2, and 5mM). Cancer cells were incubated with the drug for 48hs. On the day of staining, two washes with dH_2_O were performed before staining with 0.5% Crystal Violet solution for 20 minutes at room temperature with agitation. After staining, four washes were performed with dH_2_O, plates were inverted on filter paper and let air-dry for at least 2hs at room temperature. To quantify the crystal violet, Methanol was added, and the plates were incubated with the lids on for 20 minutes at room temperature on a bench rocker. Optical density was recorded at 570 nm with a BioTek Synergy H1 Hybrid Reader (Agilent). Average OD of three empty wells (background) was subtracted from the OD of each well on the plate. The average OD of non-stimulated cells was set to 100% viability and the percentage of viability for DFMO-treated cells that was calculated relative to the average OD values of cells treated with equal amount of DMSO.

### Extracellular vesicles isolation

Extracellular vesicles (EVs) were purified from 48 hours conditioned media by differential ultracentrifugation. All centrifugations were performed at 4 °C. Cell contamination was removed by centrifugation at 300 g for 10 minutes. Supernatant was then filtered using a 0.2 μm polyethersulfone filter, and an aliquot was taken to be used as a complete conditioned media. To remove apoptotic bodies and large cell debris, the supernatants were centrifuged at 3,000 g for 20 minutes. Finally, EVs were collected by ultracentrifugation in 4 or 31 mL ultracentrifugation tubes (Beckman Coulter, #355645 and #355631) at 100,000 g for 120 minutes. EVs were then washed in PBS, pelleted again for 120 minutes at 100,000 g in 50.4Ti fixed-angle rotors in a Beckman Coulter Optima XE, an resuspended for further analysis or stored at -80 °C until further use. EV size and quantification were measured using the NS300 nanoparticle characterization system (NanoSight, Malvern Instruments) equipped with a green laser (532 nm). The assays were performed according to the recommended protocol by the manufacturer. Briefly, three independent replicates of diluted EV preparations in PBS were injected at a constant rate into the tracking chamber by the provided syringe pump. The specimens were tracked at room temperature for 60 seconds. Shutter and gain were manually adjusted for optimal detection and were kept at optimized settings for all samples. An aliquot of each EV isolation was analyzed by Western Blot using anti-CD9 and anti-Tumor Susceptibility TSG101 (GenuIN Biotech, Ab#1359 and Ab#2707, respectively). Purified EVs were labeled with Vybrant DiO Cell labeling solution (Thermo Fisher, #V22886) per manufacturer’s instructions.

### Lipolysis assay

Primary white adipocytes were isolated, cultured, and differentiated as described previously (*65*). On day 6 of adipocyte differentiation, the culture media were replaced by conditioned media or the desired stimuli in DMEM supplemented with 2 % fatty acid-free BSA. After 24 hours of incubation, glycerol release was measured using Free Glycerol and Triglyceride Reagent (Sigma-Aldrich, F6428 and T2449, respectively) according to the recommended protocol by the manufacturer. Isoproterenol (Sigma-Aldrich, #1351005) at 10 μM was used as a positive control for lipolysis, while Atglistatin (Sigma-Aldrich, #SML1075) and CAY10499 (Cayman Chemical, #10007875) at 10 μM were used as negative controls. The average of either technical triplicates or quintuplicates was calculated and normalized to the non-stimulated condition. Right after cells were lysed in 0.5 % sodium deoxycholate, proteins were quantified using the BioRad DC protein assay kit (BioRad, #5000112) and stored at -80 °C until further use.

### Chemical inhibition of hypusination

To inhibit hypusination both *in vitro* and *ex vivo*, primary adipocytes or adipose tissue explants were treated with 20 µM GC7 (N1-guanyl-1,7- diaminoheptane, Sigma Aldrich, # 259545) a competitive inhibitor of DHPS. GC7 was protected from amine oxidases by the addition of 1 mM aminoguanidine (Cayman Chemical, # 81530) (*66*).

### Protein extraction, quantification, and western blotting

Snap-frozen tissues were transferred to a metal plate on wet ice and cut into pieces with a scalpel. All pieces were placed into 1.5 mL round-bottom Eppendorf tubes preloaded with 1.0 mm zirconia beads (BioSpec Products, # 11079110ZX) on wet ice. Tubes were then filled up to 500 μL of 0.5 % sodium deoxycholate solution and tissues were mechanically disrupted for 5 minutes using a Tissue Lyser II (Qiagen, #85300). Tissue homogenates were then centrifuged at 4 °C for 15 minutes at 18,500 g and the supernatants were transferred into new tubes. Cell protein lysates were prepared in 0.5% sodium deoxycholate solution and mechanically disrupted via sonication (10 pulses, 30% amplitude, 3s/2s on/off) on ice. Protein quantification was performed using BioRad DC protein assay kit (BioRad, #5000112). For Western Blot, samples were separated onto 4-12 % Bis-Tris gels (Thermo-Fisher, #NP0322BOX). Proteins were transferred to nitrocellulose membranes using the iBlot2 system (Thermo-Fisher), then treated with blocking buffer (5 % BSA in TBS-Tween 0.1 %) for 1 h at room temperature. Membranes were incubated overnight at 4 °C in primary antibody diluted in blocking buffer, washed three times for 10 min with TBS-Tween 0.1 % and then incubated in horseradish peroxidase (HRP)-conjugated secondary antibody diluted in blocking buffer for 1 h at room temperature. Membranes were washed again in TBS-Tween 0.1 % as described above, and signal was revealed with Pico ECL reagent and imaged using ImageQuant 800 (Amersham).

### RNA extraction and qPCR

mRNA was extracted from frozen tissues in TRIzol (Thermo Fisher # 15596018), purified using the Direct-Zol RNA MiniPrep plus kit (Zymo Research #R2052)per manufacturer’s instructions. Concentration and purity of aqueous mRNA was assessed using a Tak3 plate with a Synergy H1 plate reader (BioTek). Taqman-based quantitative real-time PCR was performed using the SuperScript III Platinum One-Step qRT-PCR reagent (Thermo Fisher #11732088). 20 ng RNA was loaded per reaction in a 10 μL volume, with samples run in triplicate and multiplexed to normalize to the internal housekeeping control gene 36B4. Alternatively, reverse transcription was performed using iScript cDNA Synthesis Kit (BioRad, #1708891) and qRT-PCR was performed using PowerUp SYBR Green Master Mix (Applied Biosystems, #A25742). All samples were run in triplicate and the ddCT values were normalized to calculate relative expression of each gene. The fold expression was calculated using the ΔΔCt method with 36B4 as a reference gene. In all cases, qPCR protocol and subsequent analysis was executed using a QuantStudio 5 thermal cycler (Applied Biosystems). Primer and probe sequences are included in supplementary information (Supplementary Table S2).

### Fully Automated Artificial Intelligence-Based CT Muscle Biomarkers

The artificial intelligence (AI) CT tools were developed using approximately 44,735 training cases derived from two independent institutions: the University of Wisconsin School of Medicine and Public Health and the National Institutes of Health (NIH), as described before (*52*). The system has received commercial licensing for clinical implementation and has been independently validated at multiple sites (*49, 51, 67-78*). Each AI tool within the container operates as an independent module, and the performance of one module does not influence the functionality of others. The initial preprocessing stage standardizes the image slice thickness and spacing to 3 mm and applies corrections for CT attenuation offsets. Following this, a body-part regression algorithm assesses the adequacy of anatomic coverage and identifies the positions of key landmarks (*79, 80*). Subsequent automated segmentation generates quantitative metrics for cross-sectional area (cm^2^) and mean attenuation (HU) of skeletal muscle at L3 and cross-sectional areas and attenuations for VAT, SAT, TAT (=VAT + SAT) and IMAT at L3. Segmentation maps were also generated for visual quality checks. Reference ranges for each biomarker, as previously established as done before (*51, 74, 77, 81*). Examinations were excluded if the patient position was non-supine (e.g., prone or decubitus), determined by DICOM header information or body-part regression output; if anatomic coverage was incomplete, such as missing the region from the top of the diaphragm through the lower poles of the kidneys and vertebral levels L1-L3, or if technical limitations such as restricted field of view or significant artifact prevented successful AI processing. The criteria and automated exclusion procedures were consistent with those described in prior work (*72, 82*).

### Human serum sample assays

Samples were collected between 2012 and 2020. Patients were in a fasted state. Samples were processed within 4 hours after collection. Blood samples were centrifuged at room temperature for 15 minutes at 1500 x g and the separated plasma was stored at -80 °C until assayed at the University of Wisconsin Madison Carbone Cancer Center Pharmacology Laboratory. GDF-15 (kit # K151YDR-2) concentrations were measured using the respective Meso Scale Discovery (MSD) R-PLEX electrochemiluminescent immunoassay kits. Plasma IL-6 concentrations were measured using the MSD U-PLEX assay kit (K151TXK-1). Standards and samples were applied in duplicate wells in the assay plates and read in a MESO QuickPlex SQ 120MM instrument (Rockville, MD). Plasma concentrations of muscle creatine kinase (CKM) (kit # ab264617) and free fatty acids (kit # ab65341) were measured in duplicate using sandwich ELISA immunoassay kits from Abcam (Waltham, MA). The ELISA plates were read at 450nm and 570nm, respectively.

### Synthesis of ^13^C_2_-dansyl chloride

^13^C_2_-dimethyl sulfate, phosphorous pentachloride (PCl_5_), Chloroform, dansyl chloride, L-proline, acetone, hexanes, and ethyl acetate were purchased from Sigma-Aldrich. Optima grade water, acetonitrile, formic acid, ACS grade water and methanol, 5-aminonapthalene-1-sulfonic acid, hydrochloric acid, and high glucose DMEM were purchased from Thermo Fisher Scientific. Sodium bicarbonate was purchased from Santa Cruz Biotechnology and sodium carbonate was purchased from DOT Scientific. Polyamine standards: N1-acetylspermidine was purchased from Cayman Chemical. Spermidine, Spermine, Putrescine, and N-Acetylputrescine, were purchased from Sigma-Aldrich.

A 25 mL round-bottom flask was charged with a magnetic stir bar and 3.5 mL of deionized water. 1.09 grams of sodium bicarbonate was dissolved in the flask, followed by 0.78 grams of 5-aminonapthalene-1-sulfonic acid in small portions under stirring until fully suspended. The solution was cooled in an ice bath. Once cooled, 0.77 mL of ^13^C_2_-dimethyl sulfate was added dropwise over the course of 30 minutes with continuous stirring. Reaction temperature was made to remain under 5 °C during addition. The round-bottom flask was then removed from the ice bath and heated in a water bath to 80 °C for 30 minutes with constant stirring. After heating, the reaction was allowed to cool to room temperature, and the pH was adjusted to 4.0 using 0.46 mL concentrated HCl. A yellow precipitate was collected by vacuum filtration, washed with cold water, and allowed to air-dry to a constant weight. This dimethylamino intermediate was then further dried at 120 °C for 1 hour. The yellow precipitate was ground under ice-water cooling into a fine powder and transferred to a dry flask. 0.88 grams of PCl_5_ were added and flask contents thoroughly mixed. The mixture was heated to 60 °C in strict moisture-free conditions for 2 hours. After reaction, the contents of the flask were allowed to cool to room temperature, and the contents were quenched with 12.5 mL of ice-cold water added dropwise. This addition was carried out with vigorous stirring in a fume hood. The aqueous mixture was neutralized slowly with careful addition of 1.75 g sodium bicarbonate in small portions. pH was kept slightly acidic to avoid hydrolysis of sulfonyl chloride. The product was extracted with 4 × 6 mL volumes of ethyl acetate, with organic layers combined immediately. The combined organic phase was then dried over anhydrous sodium sulfate, filtered, and concentrated via rotary evaporator (Buchi Rotavapor R-114). Crude product was purified via silica gel chromatography using hexane to pre-equilibrate and ethyl acetate:hexane (1:4) to elute. Fractions were monitored with thin-layer chromatography using the same running solvent and visualized under UV at 254 nm. The desired product exhibits yellow, UV-active band with an R_f_ of approximately 0.30. Purified product was dried to completion using a Speed Vac (Savant SC110) and stored in an amber vial at 4 °C. The final ^13^C_2_-dansyl chloride was obtained in a mass of 500 mg and confirmed using a Bruker Rapiflex MALDI-TOF mass spectrometer (Bruker Corp., Billerica, MA) representing a 53 % yield from 3.50 mmol of starting material

### Sample extraction for polyamine quantification

It has been reported that utilizing a classic Folch protocol (chloroform/ methanol extraction) for polyamine analysis in human serum results in lower matrix effects than other protein removal methods, presumably due to lipid depletion (*83*). This was confirmed for human plasma in this study and was selected as the extraction method of choice. Human plasma samples were stored at -80 °C until analysis. Aliquots were slowly thawed on ice and subsequently homogenized. A 100 µL aliquot of homogenous plasma was added to a clean 1.6 mL microcentrifuge tube using a positive-displacement pipette. 170 µL of ice-cold methanol and 340 µL of ice-cold chloroform were added to the plasma. This mixture was vortexed until well-incorporated, then centrifuged in an Eppendorf 5417R centrifuge (Eppendorf. Hamburg, Germany) at 16,000 g for 10 minutes at 4°C. 130 µL of the aqueous upper layer were removed and added to a clean microcentrifuge tube, with care taken to avoid disturbing the precipitate pancake.

Conditioned media were collected as previously described. A 100 µL aliquot of conditioned media was added to a clean 1.6 mL microcentrifuge tube. Proteins were precipitated by addition of 250 µL ice-cold methanol. Conditioned media samples were centrifuged at 16,000 g for 10 minutes at 4 °C, and 300 µL of supernatant were transferred to a clean microcentrifuge tube without disturbing the precipitated protein pellet.

### Sample dansylation and LC-MS/MS analysis

Calibration curve points were prepared by adding 130 µL solutions of polyamine standards in 60% methanol to clean 1.6 mL microcentrifuge tubes. Seven calibration points were generated, along with one quality control (QC) point that was later be divided into three aliquots and injected throughout the mass spectrometry run. Initial concentrations of standards in the highest calibration point were 0.075 µg/mL N-acetyl putrescine, 0.01 µg/mL N1,N12-diacetyl spermine, 0.07 µg/mL N1-acetyl spermidine, 0.14 µg/mL putrescine, 0.045 µg/mL spermidine, and 0.075 µg/mL spermine. Other calibration curve points were prepared at relative concentrations of 0.75, 0.5, 0.25, 0.1, 0.04, and 0.01 with respect to the highest calibration point for all analytes. To each sample extract and calibration point, 100 µL of 0.2 M sodium bicarbonate buffer (pH 9) was added and mixed thoroughly. Then, 50 µL of 20 mg/mL dansyl chloride in acetone was added. Tubes were capped and allowed to react for 6 hours at room temperature in the dark. After incubation, 50 µL of aqueous 200 mg/mL L-proline was added to quench the reaction for 15 minutes at room temperature in the dark. For preparation of ^13^C_2_-labeled internal standards, 150 µL of polyamine standards in water were added to a clean 1.6 mL microcentrifuge tube. The standards were present in concentrations of 1.0 µg/mL N-acetyl putrescine, 0.15 µg/mL N1,N12-diacetyl spermine 1.0 µg/mL N1-acetyl spermidine, 1.8 µg/mL putrescine, 0.6 µg/mL spermidine, and 1.0 µg/mL spermine. This solution was derivatized and quenched identically to the sample extracts, except ^13^C_2_-dansyl chloride was utilized and all volumes were tripled. After quenching, the heavy-tagged internal standards were extracted with one addition of 1200 µL ethyl acetate. 300 uL of deionized water were added to encourage layer formation. The layers were fully mixed and allowed to separate, after which 1050 µL of the upper organic layer were removed and added to a clean 30 mL conical tube. This was diluted to a final volume of 21 mL in ethyl acetate and utilized in the subsequent extraction of tagged samples and calibration curve points. Each calibration curve point, extracted sample, and quality control point was extracted with 400 µL of the ethyl acetate containing internal standards produced above and 100 µL deionized water to encourage layer formation. Samples and standard points were mixed thoroughly and allowed to separate. 350 µL of the organic upper phase were removed, added to a clean microcentrifuge tube, and dried to completion in an Eppendorf Vaccufuge (Eppendorf. Hamburg, Germany). Dried samples were stored in the dark at -20 °C until reconstitution and LCMS/MS analysis.

Samples were reconstituted in 50 µL of 50/50 ACN/H_2_O, gently mixed for 30 minutes, then centrifuged at 16,000 g for 10 minutes at 4 °C. 45 µL of the supernatant were transferred to a glass total recovery LCMS vial (Waters Corp., Milford MA). Samples were reconstituted no more than 6 hours before analysis and held at 4 °C in the LC autosampler. Triplicate 7 µL injections of samples, QC points, and calibration points were separated and analyzed on a Waters Acquity UPLC system (Waters Corp.) coupled to a SCIEX QTRAP 5500 mass spectrometer operating in positive-ion mode (AB SCIEX LLC, Marlborough MA). A 100 mm x 2.1 mm Kinetix 2.6 µm C18 column (Phenomenex Inc., Torrance CA) with attached C18 guard column was utilized for separation. A flow rate of 450 µL/min and column temperature of 25 °C was employed. Mobile phase A was 100 % Optima grade water with 0.1 % formic acid, and mobile phase B was 100 % Optima grade acetonitrile with 0.1 % formic acid. A gradient flow profile was employed, starting with 1 % mobile phase B from 0 minutes to 2 minutes, with a linear increase to 40 % B at 7.5 minutes. % B was kept constant from 7.5 to 9 minutes and increased linearly to 70 % at 15 minutes and again to 98 % at 17.25 minutes. % B was held at 98 % until minute 19, where it was ramped back down to 1 % between minutes 19-20. The flow was held constant for one more minute to allow for column equilibration before the subsequent injection. An internal divert valve was used to direct the flow path to waste for the first minute of analysis. MRM source and transition parameters (Supplementary Table S3) were optimized via direct infusion of standards and scheduled based on compound retention times. Method development and optimization were performed using SCIEX Analyst control software, and Skyline MS was utilized for data processing and quantitation (*84*). Linear or quadratic regression was selected for fitting on an analyte-by-analyte basis due to varying dynamic ranges and instrument responses across the analytes being profiled. Certain calibration points were discarded at the low or high end of calibration curves on an analyte-by-analyte basis if they fell significantly far outside the instrument’s usable response range, and only if they did not overlap with any unknown sample signals. No interior calibration points were excluded for any reason. All analyte calibration curves returned R^2^ values equal to or above 0.998. Identical aliquots of dansylated standards were prepared and injected as quality control samples at the beginning, middle, and end of the mass spectrometry run to monitor any instrument drift, and spike recovery analyses were conducted to ensure acceptable matrix effects.

### Statistical analysis

Cell culture data are presented as mean ± SD. Data from mouse studies are presented as mean ± SEM. Statistical analysis was performed on Prism software (GraphPad) using Student’s t-test for comparisons between two groups, one-way ANOVA with multiple comparisons for assessment of more than two groups, and two-way ANOVA with multiple comparisons for repeated measurements. Comparisons among specific groups were done using post-tests and are indicated in the figure legends. The statistical parameters (i.e., n number and p values) are reported in the figure legends.

## ACKNOWLEDGMENTS

We thank Drs. Dave Harris, Valentina Lo Sardo, Snehal Chaudhari, Vincent Cryns, Richard Anderson, Noelle LoConte, and Jason Cantor (University of Wisconsin-Madison) for valuable discussions and comments on the manuscript; Drs. Marcelo Vargas and Mariana Bresque (University of Wisconsin-Madison SMPH) for technical assistance with immunofluorescence staining; the Head and Neck Cancer SPORE, the Translational Science Biocore, and the Pharmacology Laboratory, supported by P30 CA014520, at UW Carbone Cancer Center for providing the H&N cancer cell lines and for use of its facilities and services including technical assistance with serum samples from newly diagnosed PDAC patients. AI-assisted proofreading was used to correct grammar and style. The models in Figures 1, 2, S1, and S2 were created using BioRender.com.

## AUTHOR CONTRIBUTIONS

Conceptualization: AG, AJK.

Methodology: MF, KO, CMA, JWG, PJP, JJC, CAL, MPdM, LL, AJK, AG.

Investigation: MF, RJF, JKH, CJK, JL, AN, EJS, KK, NR, AJ, LC.

Visualization: MF, AG.

Funding acquisition: AG, AJK, LL, MF, CJK.

Writing original draft: MF, AG.

Writing, review, and editing: MF, RJF, JKH, CJK, JL, AN, EJS, KK, NR, AJ, LC, KO, CMA, JWG, PJP, JJC, CAL, MPdM, LL, AJK, AG.

## DATA AND MATERIALS AVAILABILITY

No new biomaterials or reagents were generated in this study. Any requests for reagents and protocols can be directed to AJK or AG.

## FIGURES

**Figure S1.**
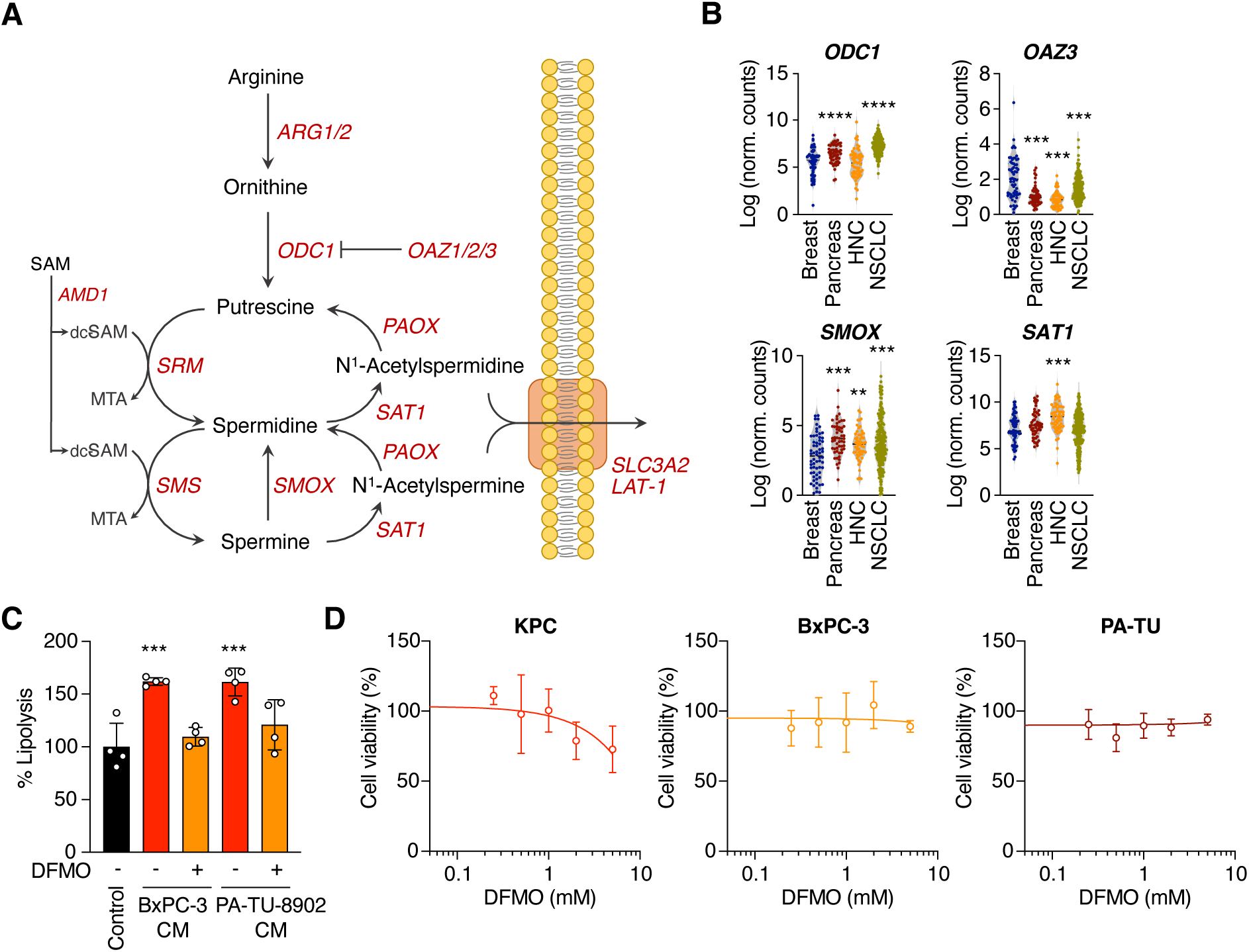
The polyamine biosynthetic pathway is dysregulated in highly cachectic cancers. (**A**) Schematic representation of the polyamine biosynthetic pathway. (**B**) ODC1, OAZ3, SMOX, and SAT1 are significantly higher in cancer cells derived from highly cachectic cancers compared to cells obtained from less chachectic tumors (i.e., breast). Data from Li et al., 2019 (*29*). (**C**) Inhibition of polyamine synthesis with 1 mM DMFO blocks the pro-lipolytic effects of BxPC3 and PA-TU-8902 CM (n = 4 per condition). (**D**) Dose-dependent effect of DFMO on cancer cell viability. Data shown ad mean ± SD (D). **p<0.01, ***p<0.001, 1-way ANOVA with Tukey’s (B) or Dunnett’s (E) post-test.

**Figure S2.**
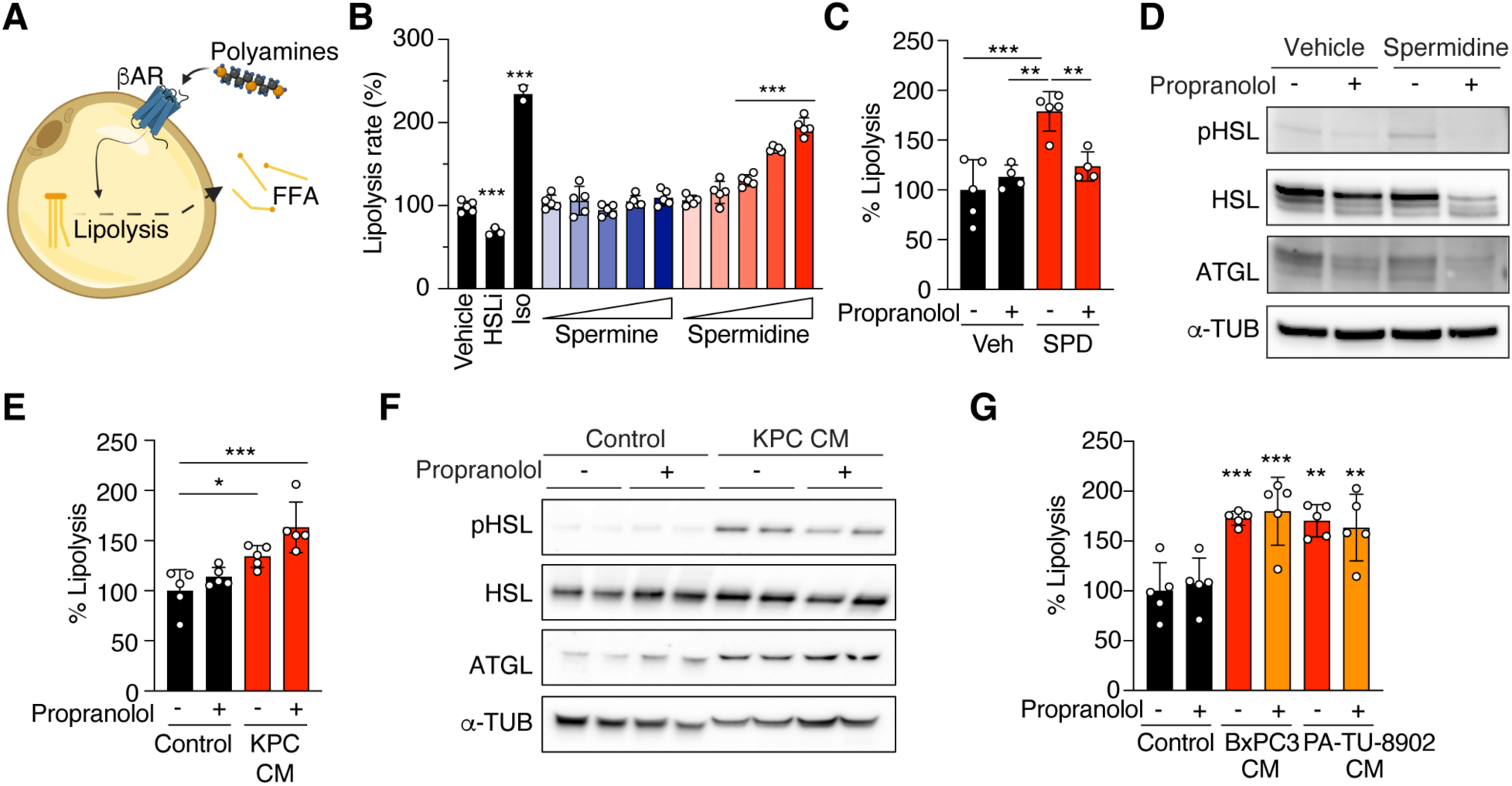
Cancer-derived polyamines do not promote lipolysis via activation of the adrenergic receptor. (**A**) Free polyamines may act as ligands for the adrenergic receptor. (**B**) Polyamines activate lipolysis in a dose- and species-specific manner. (**C** and **D**) Inhibition of the adrenergic receptor with propranolol prevents spermidine-induced activation of lipolysis. (**E**, **F**) Propranolol does not prevent KPC CM-induced activation of lipolysis. (**G**) Similar to KPC CM, the activation of lipolysis in response to Bx-PC3 and PA-TU-8902 CM is not blocked by co-treatment with propranolol. (**H**) Representative NS300 nanoparticle system profiles of three independent isolation of KPC EVs. (**I**) EV markers TSG101 and CD9, but not hypusine or eIF5A, can be detected in purified EVs. (**J**) Representative images of fluorescently labeled EVs taken up by adipocytes 24 hours post-incubation. Data shown ad mean ± SD. *p<0.05, **p<0.01, ***p<0.001, 1-way ANOVA with Dunnett’s post-test.

**Figure S3.**
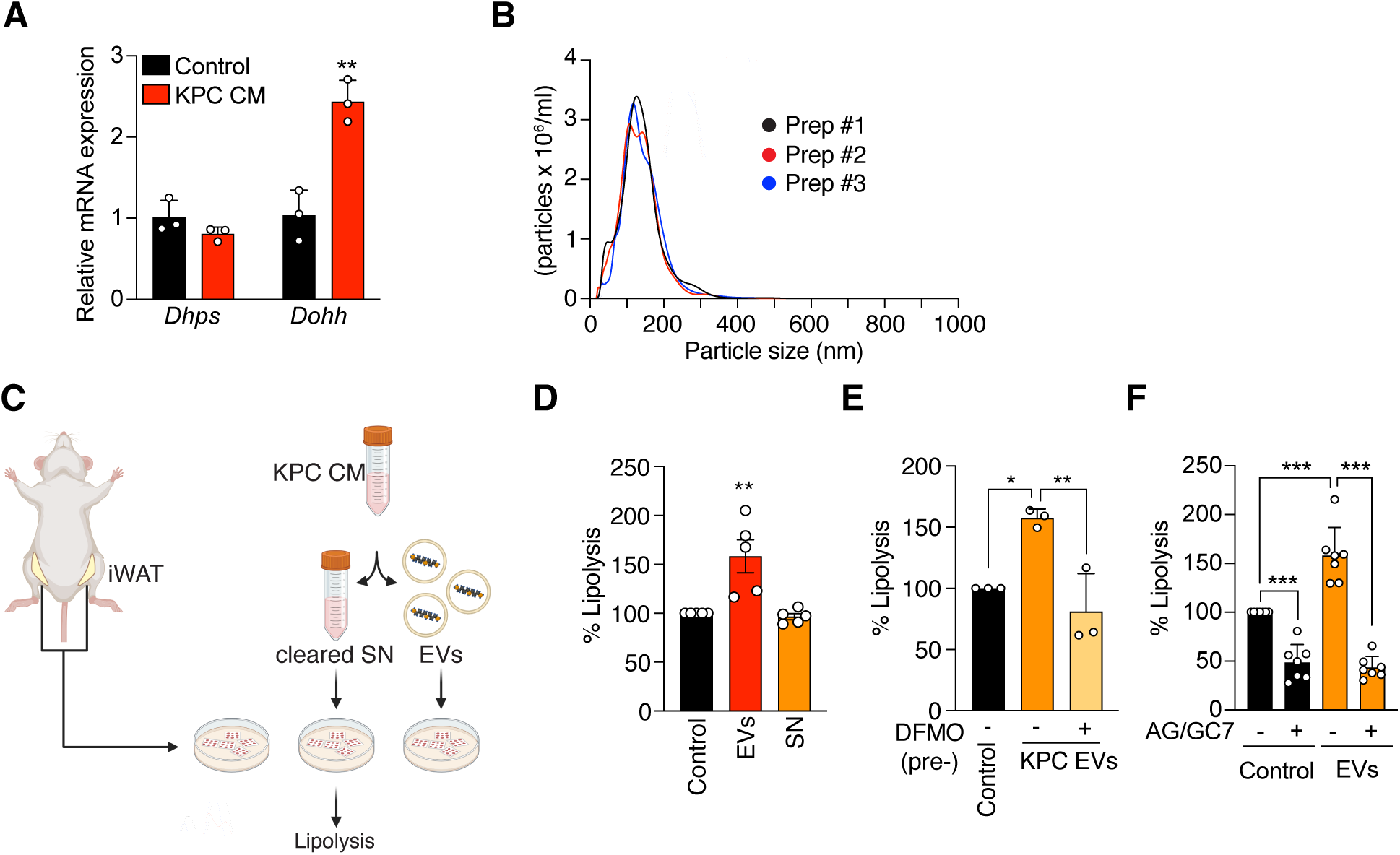
Cancer-derived polyamines are delivered to the adipocytes via extracellular vesicles. (**A**) Expression levels of *Dhps* and *Dooh* in adipocytes exposed to KPC CM for 24 hours (n = 3). (**B**) Representative NS300 nanoparticle system profiles of three independent isolation of KPC EVs. (**C**) Experimental design of *ex vivo* assessment of tumor-adipose crosstalk. (**D**) KPC EVs, but not EV-depleted KPC CM promote lipolysis in iWAT *ex vivo* (n = 5). (**E**) Inhibition of polyamine synthesis in KPC cells abolishes the pro-lipolytic effects of KPC EVs (n = 3). (**F**) Similarly, treatment of iWAT explants with the hypusination inhibitor GC7 blocks EVs’ lipolytic effects (n = 7). Data shown ad mean ± SD. *p<0.05, **p<0.01, ***p<0.001, unpaired t test (A) or 1-way ANOVA with Dunnett’s post-test (D, E, and F).

**Figure S4.**
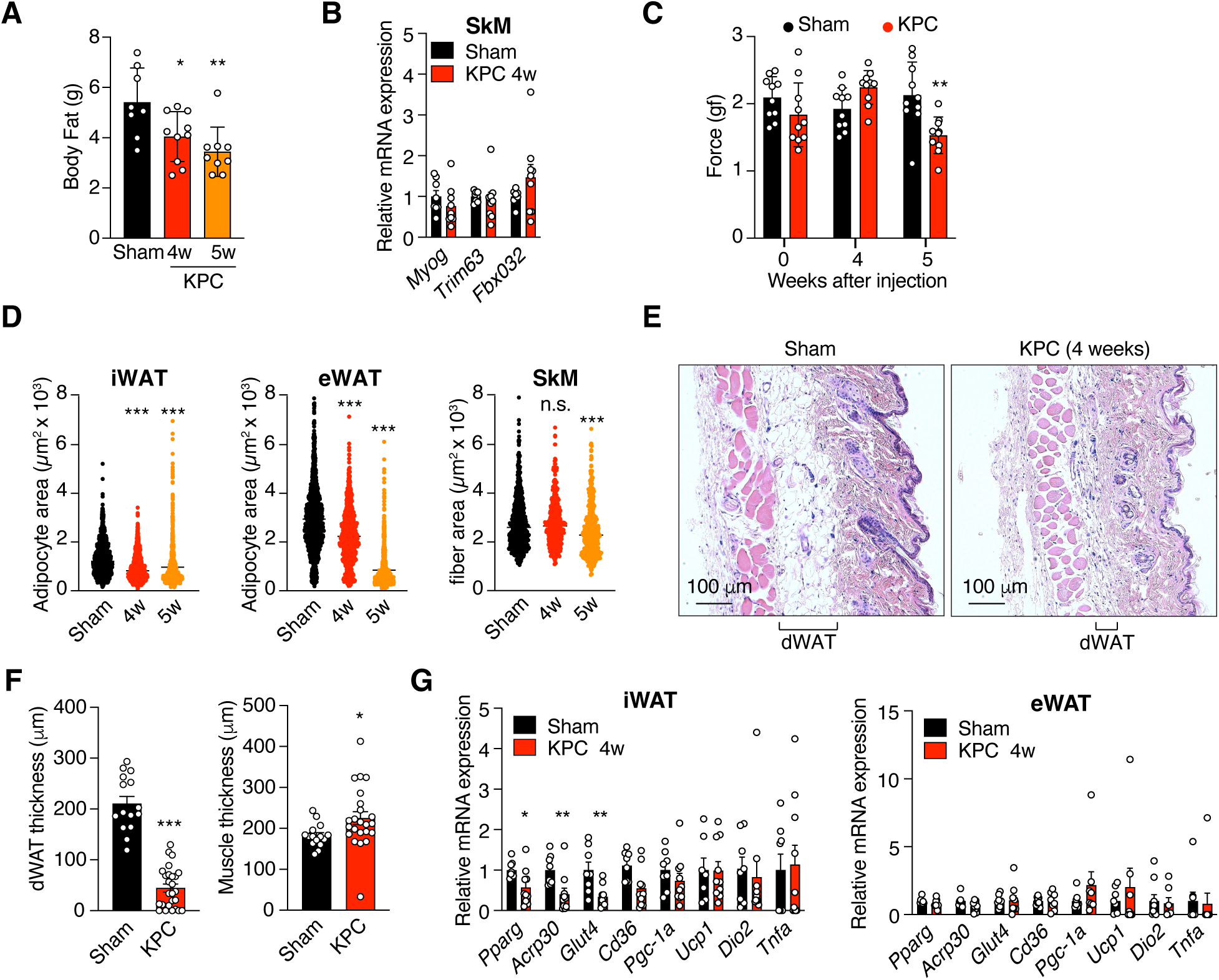
Adipose tissue metabolic rewiring precedes skeletal muscle atrophy and inflammation in KPC tumor-bearing mice. (**A**) Body fat content in sham and tumor-bearing mice at 4 and 5 weeks after KPC cells implantation (n = 8 sham, 10 KPC 4-week, 9 KPC 9-week). (**B**) Expression of atrophying markers in skeletal muscle isolated from sham and tumor-bearing mice 4 weeks post implantation (n = 8 sham, 9 KPC 4-week). (**C**) Grip strength in sham and tumor-bearing mice at baseline and after 4- and 5-weeks post implantation (n = 10 mice per group). (**D**) Quantification of adipocyte size and skeletal muscle fiber area in sham and tumor-bearing animals (n = 6 animals per condition were imaged and analyzed). (**E** and **F**) Representative images of dermal adipose tissue (dWAT) and quantification of dWAT and muscle thickness in sham and tumor-bearing mice 4 weeks after implantation (n = 14-15 sham, 23-26 KPC). (**G**) mRNA levels of classical markers of adipocytes, thermogenic in inflammatory markers in iWAT and eWAT of sham and tumor-bearing mice at 4 weeks (n = 8 sham, 9 KPC). Data shown as mean ± SEM. *p<0.05, **p<0.01, ***p<0.001, 1-way ANOVA with Dunnett’s post-test (A, D) or unpaired t test (C, F, and G).

**Figure S5.**
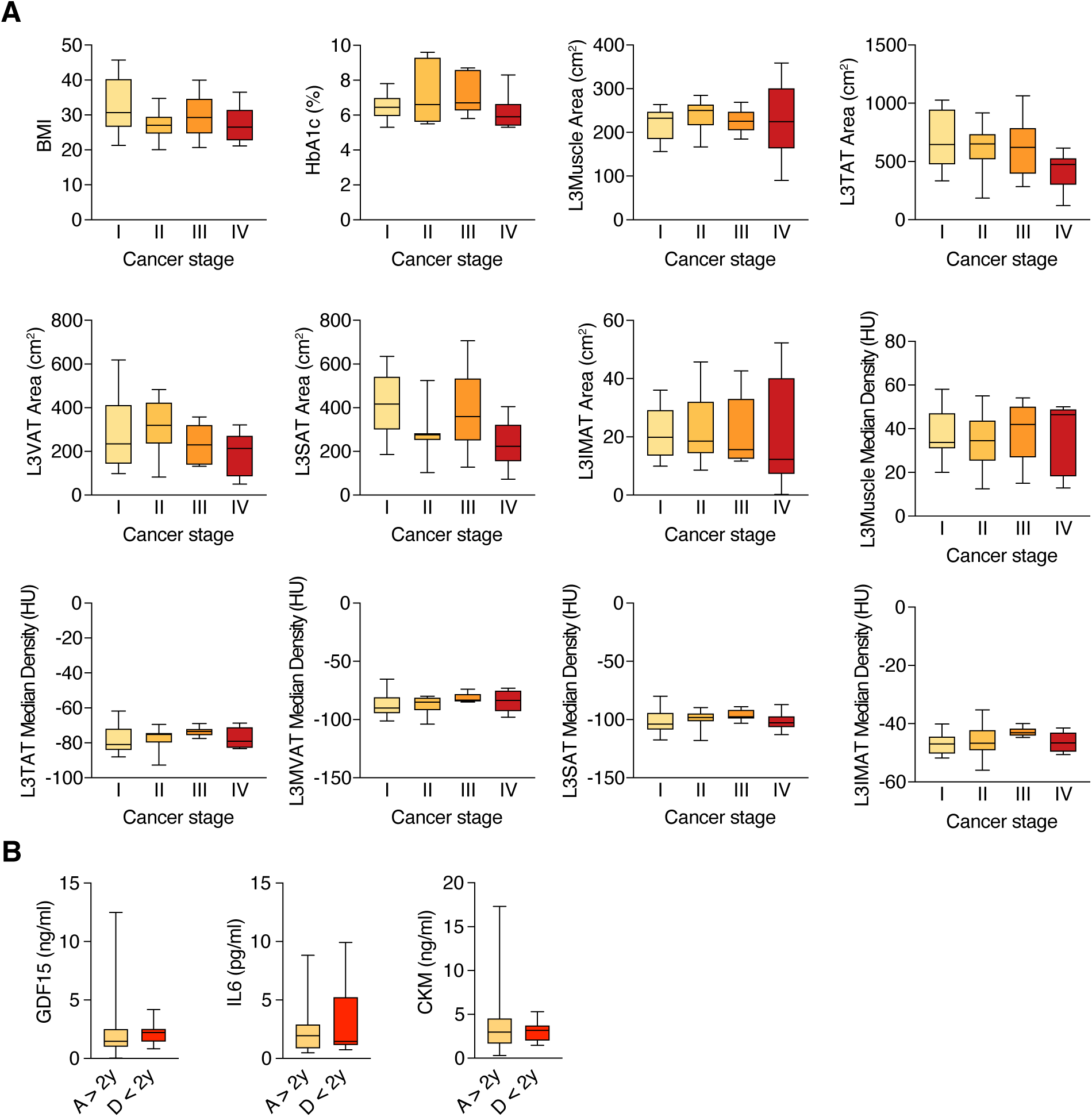
Biomarkers of advance cachexia at diagnosis. (**A**) BMI, HbA1c, and CT-derived metrics are not associated with cancer stage (n = 12 stage I, 13 stage II, 6 stage III and IV, respectively). (**B**) Circulating levels of GDF15, IL6, and muscle Creatine Kinase (CKM) at diagnosis are not predictive of survival outcomes (n = 30 A > 2y, 9 D < 2y).

**Supplementary Table 1.**
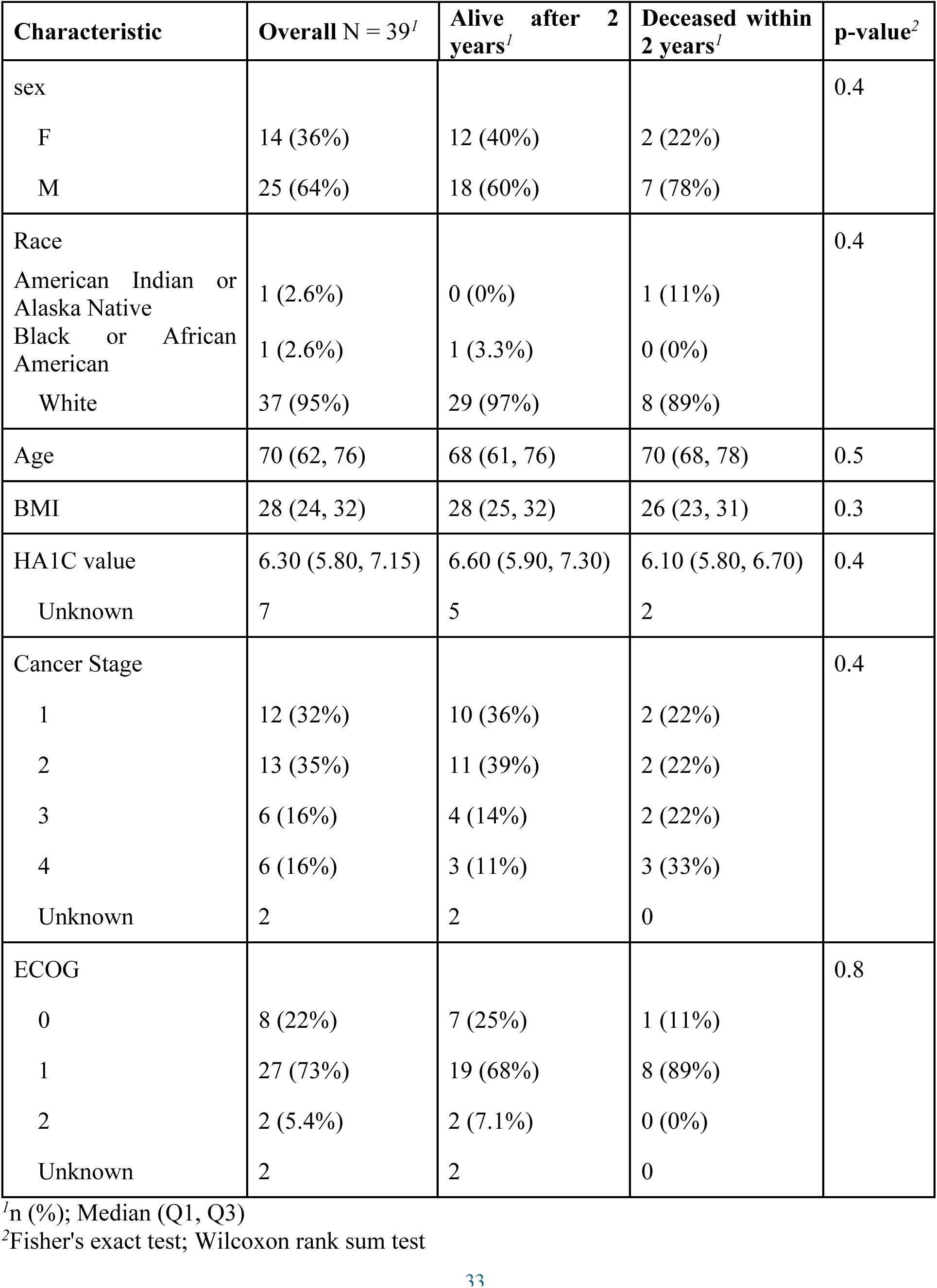
Clinical data of patients newly diagnosed with PDAC.

| Characteristic | Overall N = 39<sup>1</sup> | Alive after 2 years<sup>1</sup> | Deceased within 2 years<sup>1</sup> | p-value<sup>2</sup> |
| --- | --- | --- | --- | --- |
| sex |  |  |  | 0.4 |
| F | 14 (36%) | 12 (40%) | 2 (22%) |  |
| M | 25 (64%) | 18 (60%) | 7 (78%) |  |
| Race |  |  |  | 0.4 |
| American Indian or Alaska Native | 1 (2.6%) | 0 (0%) | 1 (11%) |  |
| Black or African American | 1 (2.6%) | 1 (3.3%) | 0 (0%) |  |
| White | 37 (95%) | 29 (97%) | 8 (89%) |  |
| Age | 70 (62, 76) | 68 (61, 76) | 70 (68, 78) | 0.5 |
| BMI | 28 (24, 32) | 28 (25, 32) | 26 (23, 31) | 0.3 |
| HA1C value | 6.30 (5.80, 7.15) | 6.60 (5.90, 7.30) | 6.10 (5.80, 6.70) | 0.4 |
| Unknown | 7 | 5 | 2 |  |
| Cancer Stage |  |  |  | 0.4 |
| 1 | 12 (32%) | 10 (36%) | 2 (22%) |  |
| 2 | 13 (35%) | 11 (39%) | 2 (22%) |  |
| 3 | 6 (16%) | 4 (14%) | 2 (22%) |  |
| 4 | 6 (16%) | 3 (11%) | 3 (33%) |  |
| Unknown | 2 | 2 | 0 |  |
| ECOG |  |  |  | 0.8 |
| 0 | 8 (22%) | 7 (25%) | 1 (11%) |  |
| 1 | 27 (73%) | 19 (68%) | 8 (89%) |  |
| 2 | 2 (5.4%) | 2 (7.1%) | 0 (0%) |  |
| Unknown | 2 | 2 | 0 |  |

<sup>1</sup>n (%); Median (Q1, Q3)
<sup>2</sup>Fisher's exact test; Wilcoxon rank sum test

**Supplementary Table 2.**
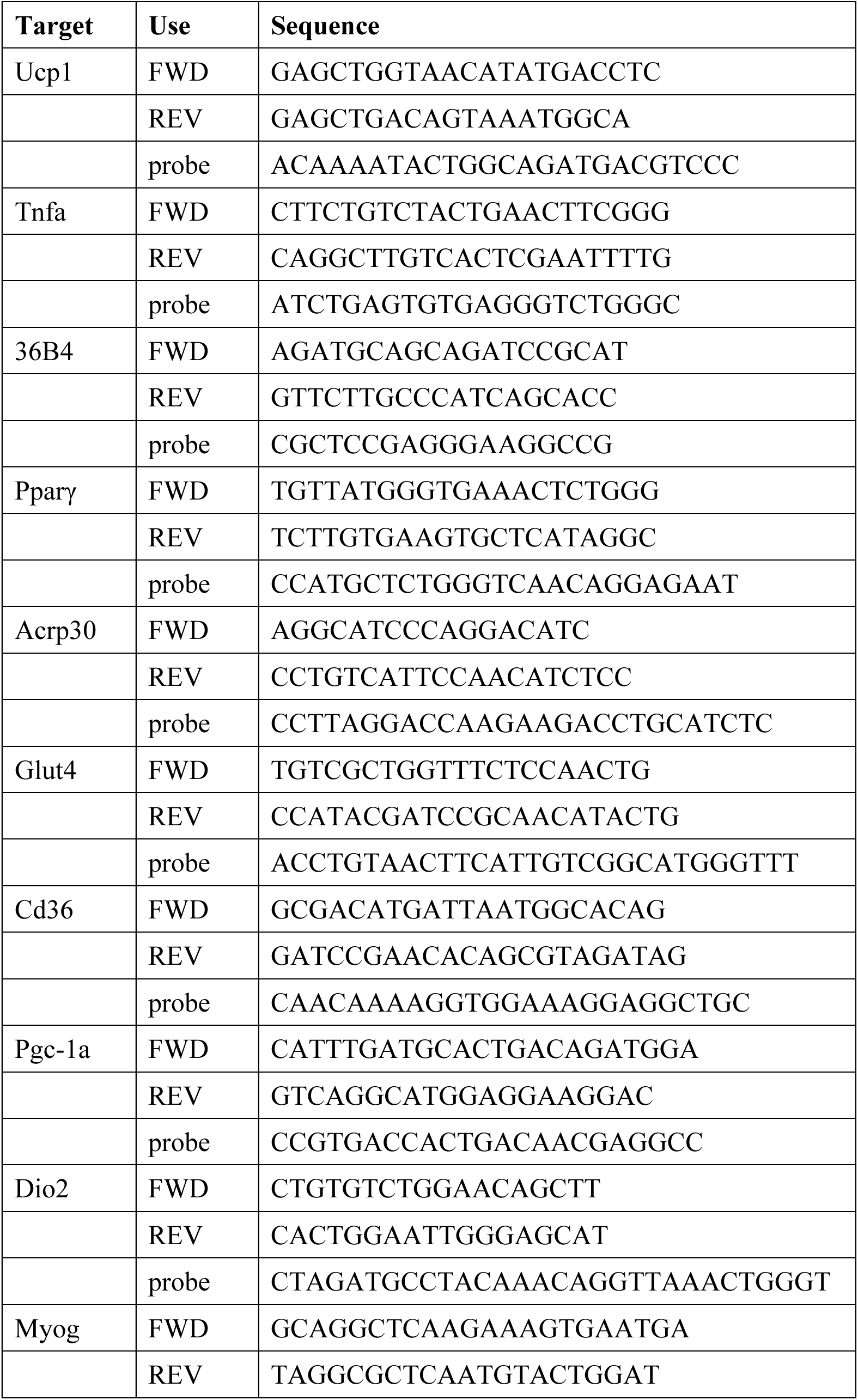
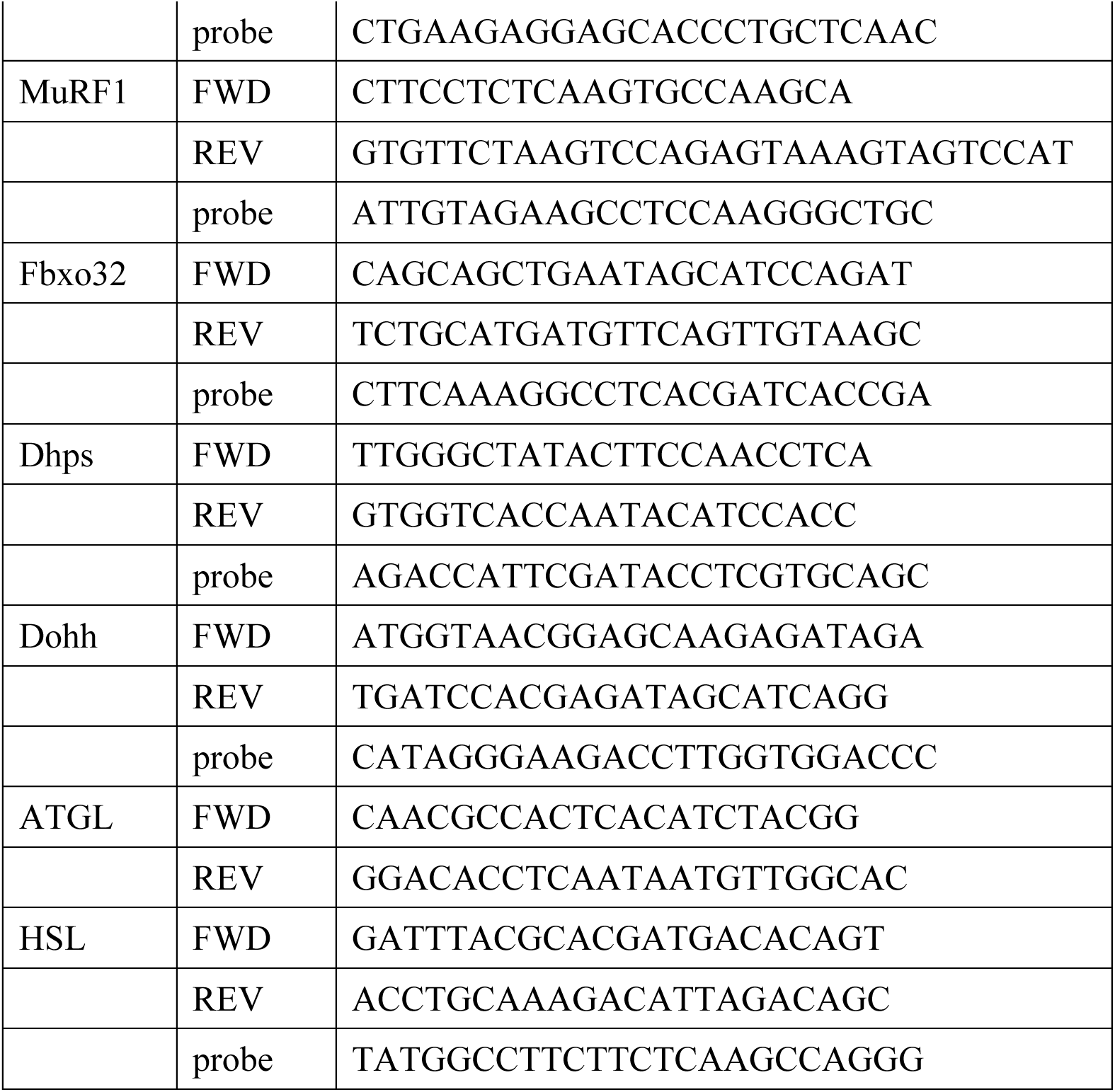
Sequences of primers and probes used in this study.

**Supplementary Table S3.** MRM mass spectrometry source and transition parameters for LC-MS/MS polyamine detection.

|  | **PUT** | **Ac-PUT** | **SPD** | **Ac-SPD** | **SPM** | **DiAc-SPM** |
| --- | --- | --- | --- | --- | --- | --- |
| Retention time (min) | 12.7 | 6.8 | 16 | 11.7 | 17.2 | 10.7 |
| MS1 m/z | 555.1 | 364.1 | 845.4 | 654.3 | 1135.6 | 753.2 |
| MRM1 m/z | 234.1 | 322.1 | 360.1 | 100.1 | 360.1 | 502.1 |
| MRM2 m/z | 220.1 | 305.1 | 303.1 | 305.1 | -- | 100.1 |
| IS MS1 m/z | 559.1 | 366.1 | 851.4 | 658.3 | 1143.6 | 757.2 |
| IS MRM1 m/z | 236.1 | 324.1 | 362.1 | 100.1 | 362.1 | 504.1 |
| IS MRM2 m/z | 222.1 | 307.1 | 305.1 | 307.1 | -- | 100.1 |
| MRM1 CE | 25 | 20 | 50 | 25 | 65 | 45 |
| MRM2 CE | 25 | 27 | 50 | 25 | -- | 99 |
| Spray V | 1500 | 2000 | 4000 | 2000 | 4500 | 3000 |
| Source temp. | 500 | 450 | 500 | 550 | 450 | 450 |

## REFERENCES

1. J. M. Argiles, S. Busquets, B. Stemmler, F. J. Lopez-Soriano, Cancer cachexia: understanding the molecular basis. Nat Rev Cancer 14, 754–762 (2014).

2. K. Fearon, J. Arends, V. Baracos, Understanding the mechanisms and treatment options in cancer cachexia. Nat Rev Clin Oncol 10, 90–99 (2013).

3. E. Bruera, C. Sweeney, Cachexia and asthenia in cancer patients. Lancet Oncol 1, 138–147 (2000).

4. S. C. Teunissen et al., Symptom prevalence in patients with incurable cancer: a systematic review. J Pain Symptom Manage 34, 94–104 (2007).

5. W. D. Dewys et al., Prognostic effect of weight loss prior to chemotherapy in cancer patients. Eastern Cooperative Oncology Group. Am J Med 69, 491–497 (1980).

6. A. G. Moses, J. Maingay, K. Sangster, K. C. Fearon, J. A. Ross, Pro-inflammatory cytokine release by peripheral blood mononuclear cells from patients with advanced pancreatic cancer: relationship to acute phase response and survival. Oncol Rep 21, 1091–1095 (2009).

7. E. J. Roeland et al., Management of Cancer Cachexia: ASCO Guideline. J Clin Oncol 38, 2438–2453 (2020).

8. M. Maltoni et al., High-dose progestins for the treatment of cancer anorexia-cachexia syndrome: a systematic review of randomised clinical trials. Ann Oncol 12, 289–300 (2001).

9. C. L. Loprinzi et al., Randomized comparison of megestrol acetate versus dexamethasone versus fluoxymesterone for the treatment of cancer anorexia/cachexia. J Clin Oncol 17, 3299–3306 (1999).

10. A. Pascual Lopez et al., Systematic review of megestrol acetate in the treatment of anorexia-cachexia syndrome. J Pain Symptom Manage 27, 360–369 (2004).

11. V. Ruiz Garcia, E. Lopez-Briz, R. Carbonell Sanchis, J. L. Gonzalvez Perales, S. Bort-Marti, Megestrol acetate for treatment of anorexia-cachexia syndrome. Cochrane Database Syst Rev, CD004310 (2013).

12. A. Jatoi et al., Dronabinol versus megestrol acetate versus combination therapy for cancer-associated anorexia: a North Central Cancer Treatment Group study. J Clin Oncol 20, 567–573 (2002).

13. A. Jatoi, Weight loss in patients with advanced cancer: effects, causes, and potential management. Curr Opin Support Palliat Care 2, 45–48 (2008).

14. V. Lai et al., Results of a pilot study of the effects of celecoxib on cancer cachexia in patients with cancer of the head, neck, and gastrointestinal tract. Head Neck 30, 67–74 (2008).

15. A. Jatoi et al., A placebo-controlled, double-blind trial of infliximab for cancer-associated weight loss in elderly and/or poor performance non-small cell lung cancer patients (N01C9). Lung Cancer 68, 234–239 (2010).

16. D. M. Breen et al., GDF-15 Neutralization Alleviates Platinum-Based Chemotherapy-Induced Emesis, Anorexia, and Weight Loss in Mice and Nonhuman Primates. Cell Metab 32, 938–950 e936 (2020).

17. K. Fearon et al., Definition and classification of cancer cachexia: an international consensus. Lancet Oncol 12, 489–495 (2011).

18. D. Blum et al., Validation of the Consensus-Definition for Cancer Cachexia and evaluation of a classification model--a study based on data from an international multicentre project (EPCRC-CSA). Ann Oncol 25, 1635–1642 (2014).

19. M. Rydén et al., Lipolysis—Not inflammation, cell death, or lipogenesis—Is involved in adipose tissue loss in cancer cachexia. Cancer 113, 1695–1704 (2008).

20. T. Agustsson et al., Mechanism of increased lipolysis in cancer cachexia. Cancer research 67, 5531–5537 (2007).

21. K. G. Burfeind, K. A. Michaelis, D. L. Marks, The central role of hypothalamic inflammation in the acute illness response and cachexia. Semin Cell Dev Biol 54, 42–52 (2016).

22. S. Kir et al., Tumour-derived PTH-related protein triggers adipose tissue browning and cancer cachexia. Nature 513, 100–104 (2014).

23. M. Petruzzelli et al., A switch from white to brown fat increases energy expenditure in cancer-associated cachexia. Cell Metab 20, 433–447 (2014).

24. S. Elattar, M. Dimri, A. Satyanarayana, The tumor secretory factor ZAG promotes white adipose tissue browning and energy wasting. FASEB J 32, 4727–4743 (2018).

25. S. K. Das et al., Adipose Triglyceride Lipase Contributes to Cancer-Associated Cachexia. Science 333, 233–238 (2011).

26. M. Fouladiun et al., Body composition and time course changes in regional distribution of fat and lean tissue in unselected cancer patients on palliative care--correlations with food intake, metabolism, exercise capacity, and hormones. Cancer 103, 2189–2198 (2005).

27. https://sites.broadinstitute.org/ccle.

28. M. Ghandi et al., Next-generation characterization of the Cancer Cell Line Encyclopedia. Nature 569, 503–508 (2019).

29. H. Li et al., The landscape of cancer cell line metabolism. Nat Med 25, 850–860 (2019).

30. K. Igarashi, K. Kashiwagi, Modulation of cellular function by polyamines. Int J Biochem Cell Biol 42, 39–51 (2010).

31. I. Novita Sari et al., Metabolism and function of polyamines in cancer progression. Cancer Lett 519, 91–104 (2021).

32. J. Li, Y. Meng, X. Wu, Y. Sun, Polyamines and related signaling pathways in cancer. Cancer Cell Int 20, 539 (2020).

33. L. M. Shantz, V. A. Levin, Regulation of ornithine decarboxylase during oncogenic transformation: mechanisms and therapeutic potential. Amino Acids 33, 213–223 (2007).

34. S. P. Nakkina et al., Differential Expression of Polyamine Pathways in Human Pancreatic Tumor Progression and Effects of Polyamine Blockade on Tumor Microenvironment. Cancers (Basel) 13, (2021).

35. E. Monelli et al., Angiocrine polyamine production regulates adiposity. Nat Metab 4, 327–343 (2022).

36. M. H. Park, E. C. Wolff, Hypusine, a polyamine-derived amino acid critical for eukaryotic translation. J Biol Chem 293, 18710–18718 (2018).

37. A. Lipton, L. M. Sheehan, G. F. Kessler, Jr., Urinary polyamine levels in human cancer. Cancer 35, 464–468 (1975).

38. Y. Asai et al., Elevated Polyamines in Saliva of Pancreatic Cancer. Cancers (Basel) 10, (2018).

39. C. Loser, U. R. Folsch, C. Paprotny, W. Creutzfeldt, Polyamine concentrations in pancreatic tissue, serum, and urine of patients with pancreatic cancer. Pancreas 5, 119–127 (1990).

40. B. C. DeFelice et al., Polyamine Metabolites as Biomarkers in Head and Neck Cancer Biofluids. Diagnostics (Basel) 12, (2022).

41. Y. Chen et al., Machine learning to identify precachexia and cachexia: a multicenter, retrospective cohort study. Supportive care in cancer : official journal of the Multinational Association of Supportive Care in Cancer 32, 630 (2024).

42. J. Geppert, M. Rohm, Cancer cachexia: biomarkers and the influence of age. Mol Oncol 18, 2070–2086 (2024).

43. Y. Jiang et al., Imaging Cancer-associated Cachexia: Utilizing Clinical Imaging Modalities for Early Diagnosis. Radiol Imaging Cancer 7, e240291 (2025).

44. J. Ni, L. Zhang, Cancer Cachexia: Definition, Staging, and Emerging Treatments. Cancer Manag Res 12, 5597–5605 (2020).

45. M. Mourtzakis et al., A practical and precise approach to quantification of body composition in cancer patients using computed tomography images acquired during routine care. Appl Physiol Nutr Metab 33, 997–1006 (2008).

46. W. G. Looijaard et al., Skeletal muscle quality as assessed by CT-derived skeletal muscle density is associated with 6-month mortality in mechanically ventilated critically ill patients. Crit Care 20, 386 (2016).

47. T. Yoo, W. D. Lo, D. C. Evans, Computed tomography measured psoas density predicts outcomes in trauma. Surgery 162, 377–384 (2017).

48. P. M. Graffy, J. Liu, S. O’Connor, R. M. Summers, P. J. Pickhardt, Automated segmentation and quantification of aortic calcification at abdominal CT: application of a deep learning-based algorithm to a longitudinal screening cohort. Abdom Radiol (NY) 44, 2921–2928 (2019).

49. P. M. Graffy, V. Sandfort, R. M. Summers, P. J. Pickhardt, Automated Liver Fat Quantification at Nonenhanced Abdominal CT for Population-based Steatosis Assessment. Radiology 293, 334–342 (2019).

50. P. J. Pickhardt et al., Simultaneous screening for osteoporosis at CT colonography: bone mineral density assessment using MDCT attenuation techniques compared with the DXA reference standard. J Bone Miner Res 26, 2194–2203 (2011).

51. S. J. Lee et al., Fully automated segmentation and quantification of visceral and subcutaneous fat at abdominal CT: application to a longitudinal adult screening cohort. Br J Radiol 91, 20170968 (2018).

52. J. W. Garrett, P. J. Pickhardt, R. M. Summers, Methodology for a fully automated pipeline of AI-based body composition tools for abdominal CT. Abdom Radiol (NY), (2025).

53. M. Ebadi et al., Higher subcutaneous adipose tissue radiodensity is associated with increased mortality in patients with cirrhosis. JHEP Rep 4, 100495 (2022).

54. N. A. Stephens et al., Intramyocellular lipid droplets increase with progression of cachexia in cancer patients. J Cachexia Sarcopenia Muscle 2, 111–117 (2011).

55. L. Patzelt et al., MRI-Determined Psoas Muscle Fat Infiltration Correlates with Severity of Weight Loss during Cancer Cachexia. Cancers (Basel) 13, (2021).

56. J. L. Brown et al., Mitochondrial degeneration precedes the development of muscle atrophy in progression of cancer cachexia in tumour-bearing mice. J Cachexia Sarcopenia Muscle 8, 926–938 (2017).

57. V. T. Samuel, K. F. Petersen, G. I. Shulman, Lipid-induced insulin resistance: unravelling the mechanism. Lancet 375, 2267–2277 (2010).

58. N. Mir, S. A. Chin, M. C. Riddell, J. L. Beaudry, Genomic and Non-Genomic Actions of Glucocorticoids on Adipose Tissue Lipid Metabolism. Int J Mol Sci 22, (2021).

59. B. Beutler, A. Cerami, Cachectin and tumour necrosis factor as two sides of the same biological coin. Nature 320, 584–588 (1986).

60. V. A. Pipitone et al., Interleukin-1beta signalling in muscle atrophy: linking inflammation, sex-specific responses and exercise in cancer cachexia. J Physiol 602, 6641–6643 (2024).

61. G. Haemmerle et al., Hormone-sensitive lipase deficiency in mice causes diglyceride accumulation in adipose tissue, muscle, and testis. J Biol Chem 277, 4806–4815 (2002).

62. G. Haemmerle et al., Defective lipolysis and altered energy metabolism in mice lacking adipose triglyceride lipase. Science 312, 734–737 (2006).

63. M. Fabregat et al., Generation and characterization of Ccdc28b mutant mice links the Bardet-Biedl associated gene with mild social behavioral phenotypes. PLoS Genet 18, e1009896 (2022).

64. Y. Zhang et al., Canonical wnt signaling is required for pancreatic carcinogenesis. Cancer research 73, 4909–4922 (2013).

65. A. Galmozzi, B. P. Kok, E. Saez, Isolation and Differentiation of Primary White and Brown Preadipocytes from Newborn Mice. J Vis Exp, (2021).

66. B. Maier et al., The unique hypusine modification of eIF5A promotes islet beta cell inflammation and dysfunction in mice. J Clin Invest 120, 2156–2170 (2010).

67. P. J. Pickhardt et al., Biological age model using explainable automated CT-based cardiometabolic biomarkers for phenotypic prediction of longevity. Nat Commun 16, 1432 (2025).

68. P. J. Pickhardt et al., Fully Automated Deep Learning Tool for Sarcopenia Assessment on CT: L1 Versus L3 Vertebral Level Muscle Measurements for Opportunistic Prediction of Adverse Clinical Outcomes. AJR Am J Roentgenol 218, 124–131 (2022).

69. R. M. Summers et al., Atherosclerotic Plaque Burden on Abdominal CT: Automated Assessment With Deep Learning on Noncontrast and Contrast-enhanced Scans. Acad Radiol 28, 1491–1499 (2021).

70. A. A. Perez, P. J. Pickhardt, D. C. Elton, V. Sandfort, R. M. Summers, Fully automated CT imaging biomarkers of bone, muscle, and fat: correcting for the effect of intravenous contrast. Abdom Radiol (NY) 46, 1229–1235 (2021).

71. P. M. Graffy et al., Automated assessment of longitudinal biomarker changes at abdominal CT: correlation with subsequent cardiovascular events in an asymptomatic adult screening cohort. Abdom Radiol (NY) 46, 2976–2984 (2021).

72. B. D. Pooler, J. W. Garrett, A. M. Southard, R. M. Summers, P. J. Pickhardt, Technical Adequacy of Fully Automated Artificial Intelligence Body Composition Tools: Assessment in a Heterogeneous Sample of External CT Examinations. AJR Am J Roentgenol, 1–9 (2023).

73. F. A. Huber et al., AI-based opportunistic quantitative image analysis of lung cancer screening CTs to reduce disparities in osteoporosis screening. Bone 186, 117176 (2024).

74. P. J. Pickhardt, et al., Improved CT-based Osteoporosis Assessment with a Fully Automated Deep Learning Tool. Radiol Artif Intell 4, e220042 (2022).

75. P. J. Pickhardt et al., Population-based opportunistic osteoporosis screening: Validation of a fully automated CT tool for assessing longitudinal BMD changes. Br J Radiol 92, 20180726 (2019).

76. P. J. Pickhardt et al., Automated Abdominal CT Imaging Biomarkers for Opportunistic Prediction of Future Major Osteoporotic Fractures in Asymptomatic Adults. Radiology 297, 64–72 (2020).

77. P. M. Graffy et al., Deep learning-based muscle segmentation and quantification at abdominal CT: application to a longitudinal adult screening cohort for sarcopenia assessment. Br J Radiol 92, 20190327 (2019).

78. J. E. Burns, J. Yao, D. Chalhoub, J. J. Chen, R. M. Summers, A Machine Learning Algorithm to Estimate Sarcopenia on Abdominal CT. Acad Radiol 27, 311–320 (2020).

79. K. Yan et al., Deep Lesion Graphs in the Wild: Relationship Learning and Organization of Significant Radiology Image Findings in a Diverse Large-scale Lesion Database. Proceedings of the IEEE Conference on Computer Vision and Pattern Recognition, 9261-9270 (2018).

80. K. Yan, L. Lu, R. M. Summers, Unsupervised body part regression via spatially self-ordering convolutional neural networks. 2018 IEEE 15th International Symposium on Biomedical Imaging (ISBI 2018), 1022-1025 (2018).

81. S. Lee, et al., Fully Automated and Explainable Liver Segmental Volume Ratio and Spleen Segmentation at CT for Diagnosing Cirrhosis. Radiol Artif Intell 4, e210268 (2022).

82. B. D. Pooler et al., CT-Based Body Composition Measures and Systemic Disease: A Population-Level Analysis Using Artificial Intelligence Tools in Over 100,000 Patients. AJR Am J Roentgenol 224, e2432216 (2025).

83. I. Samarra et al., Gender-Related Differences on Polyamine Metabolome in Liquid Biopsies by a Simple and Sensitive Two-Step Liquid-Liquid Extraction and LC-MS/MS. Biomolecules 9, (2019).

84. K. J. Adams et al., Skyline for Small Molecules: A Unifying Software Package for Quantitative Metabolomics. J Proteome Res 19, 1447–1458 (2020).

